# RHO1-2 meganuclease gene editing targets human P23H rhodopsin-induced retinitis pigmentosa to rejuvenate rods and maintain cones

**DOI:** 10.1101/2025.09.25.678625

**Authors:** Archana Jalligampala, Jacob M. Young, Jack Feist, Wei Wang, Francesca Barone, David C. Alston, James W. Fransen, Gita Jaikumar, Kautuk Kamboj, Caitlin Mooreman, Stephen Nash, Jennifer M. Noel, Gobinda Pangeni, Joseph C. Prestigiacomo, Bhubanananda Sahu, Caitlin Turner, Henry J. Kaplan, Jonathan A. Green, Kevin D. Wells, Victor V. Bartsevich, Jon E. Chatterton, Mara Davis, Kathryn S. Evans, Janel Lape, Whitney C. Lewis, Rebecca van de Beek, Kristi D. Viles, Derek Jantz, Ronald. G. Gregg, Jeff Smith, Maureen A. McCall

## Abstract

Autosomal dominant retinitis pigmentosa (adRP) is an inherited retinal dystrophy characterized by progressive vision loss and eventual blindness. The P23H mutation (proline to histidine substitution at codon 23) in the rhodopsin (RHO) gene represents the most common form of adRP in North Americans. Currently, there is no cure for P23H adRP. Genome editing targeting the mutant RHO allele, leaving a functional wildtype (WT) allele, is an attractive approach for P23H adRP, as only one copy of RHO is needed for normal retinal function. We re-engineered an I-Cre meganuclease, called RHO1-2, to target a 22bp recognition sequence encompassing the mutation responsible for the p.P23H RHO mutation. *In vitro*, RHO1-2, cuts human P23H RHO but not WT RHO. *In vivo,* we delivered scAAV5:GRK1:RHO1-2 via subretinal injection in early-stage degeneration using the only large animal model of human p.P23H RHO adRP (TgP23H pigs). We tested RHO1-2 efficacy and durability, on retinal function using full-field electroretinograms and on retinal structure using spectral domain optical coherence tomography and immunohistochemistry. We observe that RHO1-2 treatment: arrests rod photoreceptor degeneration, resurrects rod-driven retinal function that does not exist in untreated TgP23H pigs, restores mislocalized rhodopsin expression and rebuilds rod inner and outer segments (IS/OS). Rod rescue maintains cones. A year after RHO1-2 treatment, we show that TgP23H pigs use rod-driven vision to navigate a maze. Our results demonstrate that genome editing via RHO1-2 meganuclease is a viable treatment to cure human p.P23H RHO adRP. They also suggest that meganuclease-based editors can be effective for other IRDs.

**One Sentence Summary:** Engineered meganuclease, RHO1-2 is a safe and promising therapeutic genome editing approach to cure human p.P23H RHO adRP.

## INTRODUCTION

Gene therapy uses a number of strategies to cure inherited diseases(*1–3*). For autosomal recessive diseases, lacking expression of a particular protein, gene supplementation, with an engineered expression cassette, can restore wildtype (WT) gene/protein expression and cure the disease. Gene supplementation, using AAV delivery of the missing RPE65 gene, restored vision in the FDA-approved gene therapy for Leber’s congenital amaurosis (Luxturna(*4*)). This set the stage to treat other inherited retinal degenerations (IRDs). For IRDs caused by autosomal dominant negative mutant alleles, gene augmentation with a WT allele is ineffective and extra WT copies can be cytotoxic(*5, 6*). Instead, inhibition or elimination of mutant allele expression, while sparing the WT allele, is more adventageous(*7, 8*). However, strategies to reduce mutant allele expression, including antisense oligonucleotides, ribozymes, or RNA interference(*5, 9*) have had modest success, as many eliminate both the WT and mutant alleles and then must add back a WT allele modified to resist suppression(*10–12*). Here, success depends on control of both editing efficiency of the mutant and replacement levels for WT allele/protein expression. An advantageous alternative approach uses genome editing, with the goal of selectively correcting or eliminating the mutant allele, leaving the WT allele intact and able to restore protein expression and cellular function(*7, 8, 13*).

Here, we investigated an *in vivo* genome editing strategy for autosomal dominant retinitis pigmentosa (adRP) based on re-engineered I-Cre homing endonucleases previously used to successfully reduce serum cholesterol(*14–17*). These meganucleases, referred to as ARCUS, have the advantage of small size (920bp) and highly selective recognition of long sequences (22-24bp). We targeted the most common adRP mutation in the rhodopsin gene (*RHO*) found in North Americans, p.P23H, which causes a form of the blinding eye disease, P23H adRP. Initial, *in vitro*, experiments demonstrated that our engineered ARCUS meganuclease, referred to RHO1-2, specifically targets the p.P23H *RHO* mutation and cuts a 22-base pair recognition sequence encompassing the C to A transversion mutation responsible for the p.P23H *RHO* mutation (c.68A) and does not cut the WT allele (c.68C). *In vivo* we tested the efficacy of RHO1-2 treatment via subretinal injection in the only large animal model of P23H adRP, a transgenic pig carrying the entire human p.P23H gene (rrid-NSRRC:0017; TgP23H or Tg(*18–20*). RHO1-2 treatment: arrests rod photoreceptor degeneration, resurrects rod-driven retinal function that does not exist in untreated Tg pigs, restores mislocalized rhodopsin expression and rebuilds rod inner and outer segments (IS/OS). Rod rescue maintains cone morphology and function. A year after RHO1-2 treatment, TgP23H pigs can use rod-driven vision to navigate a maze. Our proof-of-concept results demonstrate that RHO1-2 genome editing is a viable treatment for human p.P23H *RHO* adRP and suggest that meganuclease-based editors can be an effective strategy for other IRDs.

## RESULTS

### RHO1-2 meganuclease structure and gene editing of human P23H Rhodopsin

We reengineered the ARCUS meganuclease gene editing platform(*14*), and produced RHO1-2A meganuclease. Optimization to improve target site specificity produced RHO1-2B (Fig.1A, B). RHO1-2A and B differ by six amino acids (B) that are involved in DNA recognition and should control activity and specificity for the 22bp P23H *RHO* target site (Fig. 1B, c.68C>A). RHO1-2A or B discrimination of P23H *RHO* is based on the single nucleotide difference, c.68C>A, near target site’s 3′ end and should produce indels in P23H RHO, leaving the WT *RHO* gene functional (Fig. 1C). Hereon, we refer to RHO1-2A and B meganuclease as RHO1-2A and B.

**Fig. 1.**
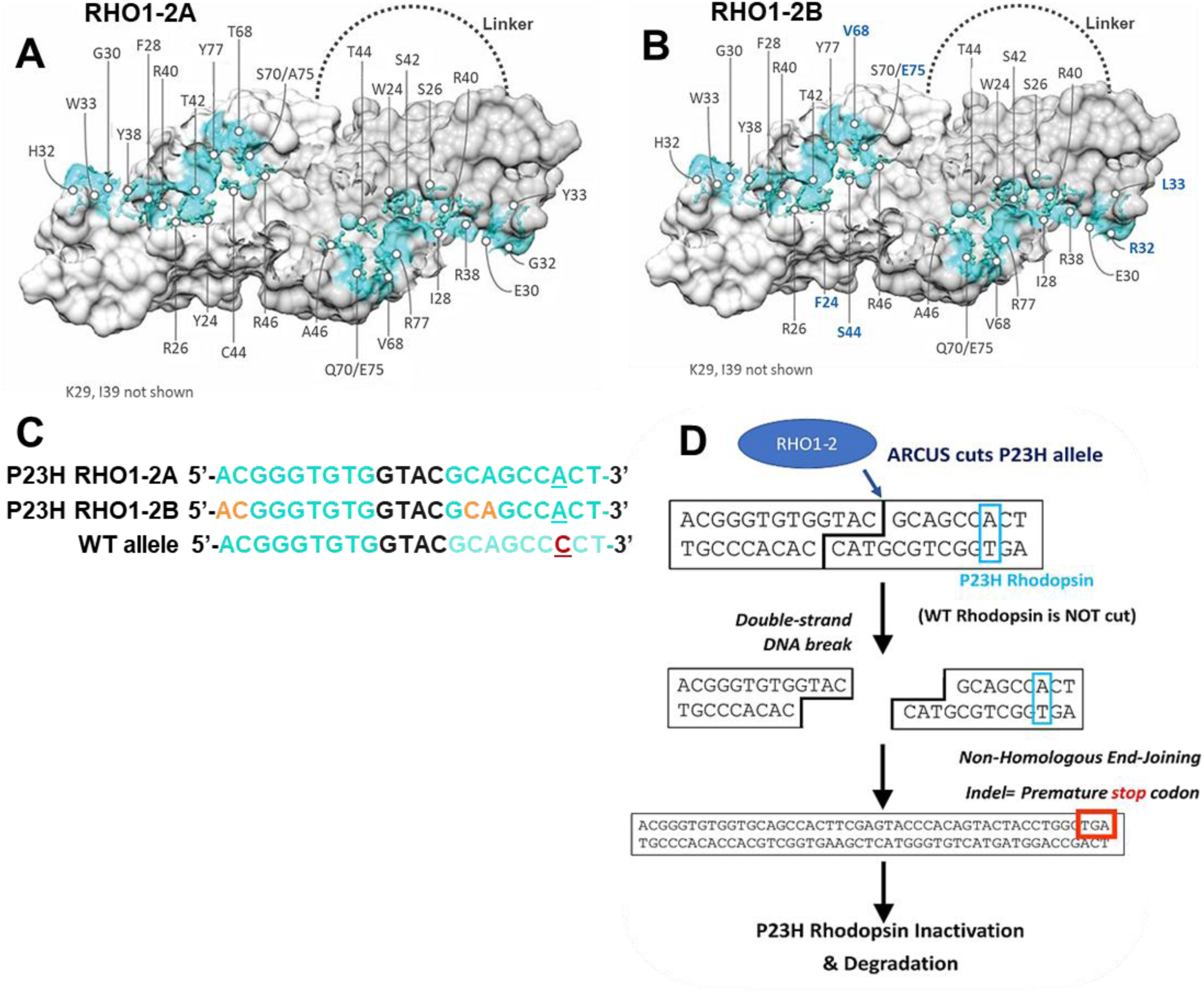
RHO1-2A & B meganuclease Structure and P23H *RHO* editing strategy. Models of the single-chain RHO1-2A (**A**) and RHO1-2B (**B**) meganucleases. The amino acids predicted to contact the meganuclease directly are shown in teal. Numbering consistent with wild-type I-CreI(*46*). (**C**) The sequence contacted by RHO1-2A and B, and the nucleotides in 1B contacted by different amino acids (orange). The c.68C>A transversion causing P23H *RHO* adRP is underlined in each. (**D**) Schematic diagram of RHO1-2 cleavage of the P23H human *RHO* target sequence, causing a double-stranded break, repaired by Non-homology End-joining (NHEJ) resulting in indels, many of which should create frameshift mutations and cause protein degradation.

### RHO1-2A and RHO1-2B distinguish between P23H and WT *RHO* DNA sequences *in vitro*

We evaluated whether RHO1-2A and B could distinguish between WT and P23H *RHO* (c.68C>A transversion) alleles in two *in vitro* cell assays. From a Flp-In CHO cell line, we created two cell lines, one carrying a 24bp fragment of the P23H human *RHO* mutant and the other the human *RHO* WT sequence (Fig. 2A, CHO-P23H and CHO-WT, respectively). Each cell line also carried a target site for Meg-23/24 (Fig. 2A, red boxes), as a positive control. We assessed cleavage in CHO-P23H and CHO-WT cells after transfection with mRNA encoding Meg23/24 and RHO1-2A or B by measuring GFP expression, and the number of GFP+ cells. RHO1-2A and B had editing efficiencies of the P23H *RHO* mutation ∼120% and 60% of the positive control, respectively (Fig. 2B). In contrast, RHO1-2A or B editing of WT target was negligible (Fig. 2C), indicating that both RHO1-2A and B efficiently cut the mutant P23H and not the WT *RHO* sequence.

**Fig. 2.**
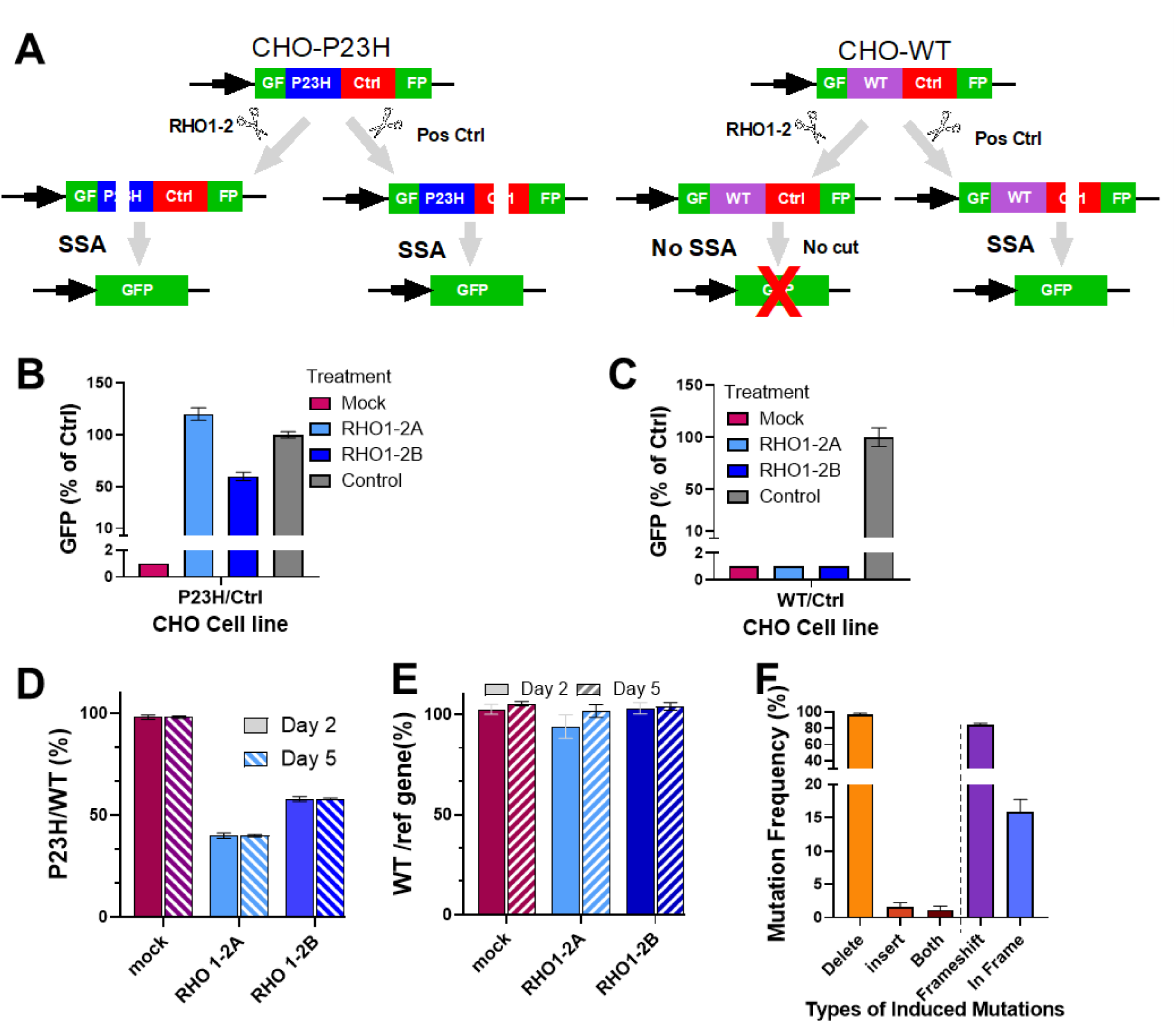
RHO1-2 meganuclease discrimination between P23H mutant and WT *RHO* alleles. (**A**) Schematic of GFP assay in CHO-P23H and CHO-WT cells. Left-Cleavage of CHO-P23H by RHO1-2A or B or the positive control meganuclease stimulates repair by single-strand annealing (SSA) and generates cells with GFP expression (GFP+). Right-RHO1-2 meganucleases do not cut CHO-WT, but the positive control does and produces GFP+ cells. GFP readout from CHO-P23H and CHO-WT cell lines. In CHO-P23H cells, (**B**) RHO1-2A and B cut the P23H target to stimulate the production of GFP+ cells. (**C**) Only positive control meganuclease generates GFP+ cells in CHO-WT. Using 2G11 cells that contain a P23H and a WT RHO allele, (**D**) the ratio of P23H to WT *RHO* mRNA is 100% in untreated (mock) and is reduced by RHO1-2 meganucleases. (**E**) The ratio of WT to a reference gene is constant, indicating that RHO1-2 meganucleases do not cut the WT *RHO* allele. (**F**) Ampseq analyses at the P23H *RHO* locus shows most mutations are indels and the majority cause a frameshift.

To determine the specificity of RHO1-2 for P23H in the presence of WT *RHO*, we generated a K562 leukemic human cell line (2G11) that carried both WT and P23H *RHO* alleles. In 2G11 cells treated with RHO1-2A or B, the number of positive droplets for the P23H (FAM) decreased relative to the WT probe (VIC; Fig. 2D). Whereas the number of FAM and VIC-positive droplets was equal in untreated cells (mock) (Fig. 2D). To ensure nucleases did not cut the WT allele, the WT probe counts were compared to a third probe on a non-targeted reference gene (Fig. 2E) . Both RHO1-2A or B produced a decrease in the P23H mutant allele copy number to ∼50% relative to WT, indicating this fraction of the cells contained indels in the P23H allele. Neither RHO1-2A nor B produced a decrease in the WT allele, the copy number compared to an unrelated reference gene was unchanged (Fig. 2E). These results further confirm that RHO1-2A and B are specific for the P23H allele and show that they have similar specificity and efficacy.

We evaluated the types of mutations induced at the cut site to determine their potential to eliminate P23H rhodopsin protein expression. Among the mutations, RHO1-2 cleavage resulted primarily in deletions in the P23H allele (Fig. 2F, orange bars) and the deletions were primarily frameshifts, predicted to eliminate P23H *RHO* expression. The remainder produced missense mutations (Fig. 2F), which also are likely to eliminate P23H expression, indicating RHO1-2 should be clinically relevant.

To determine the risk of off-target cutting beyond WT *RHO*, we evaluated genome-wide specificity of RHO1-2A and B with GUIDE-Seq, an unbiased off-target identification method adapted to meganucleases(*21*). An excess of RHO1-2A or B mRNA (1000 ng/1x10^6^ cells) and an oligodeoxynucleotide (ODN) were transfected into 2G11 cells. When ODN integration sites were identified, only sites that reoccurred in at least two of three replicates were considered potential off targets, which reduced background due to random ODN integration. For RHO1-2A, only 38 potential sites were nominated and none included WT *RHO*. Next Generation Sequencing (NGS) and an rhAmpseq panel, encompassing the 38 nominated sites, including the P23H target (Table S1), confirmed 10/38 potential sites for RHO1-2A and only 1 site for RHO1-2B, which was different from the sites verified for RHO1-2A. Further, the single off-target site verified for RHO1-2B (Chr5:4203059-4203080) was not concerning: (1) it was located in an intergenic region more than 160 kb from any coding gene, and (2) it had very low editing even with excessive nuclease input concentrations. Therefore, RHO1-2B lower off-targeting, making it the lead meganuclease for P23H adRP.

Prior to evaluation of RHO1-2A and B efficacy *in vivo*, we examined expression in WT pig retina after subretinal injection of self-complementary AAV (scAAV) to express RHO1-2A or B, spiked with GFP, all under the control of a GRK1 promoter (scAAV5:GRK1:RHO1-2A, B, or GFP (Fig. S1A). Results with RHO1-2A and B were identical. Fluorescent fundus images detected GFP expression at ≥30dpi, which remained unchanged through 300dpi. Immunohistochemical (IHC) analysis showed RHO1-2/GFP expression was limited to WT rods (Fig. S1B). In TgP23H rods, RHO1-2 and GFP expression overlapped throughout the subretinal bleb, and no expression occurred outside its margins (the transition zone, Fig. S1C and D). Within the transduced area, many rods co-expressed RHO1-2/GFP and others expressed only RHO1-2 or GFP (Fig.S1E). Areas with overlapping RHO1-2/GFP expression are referred to as RHO1-2**^+^** and those without as RHO1-2^-^.

We compared the efficacy of RHO1-2A and B *in vitro*, after subretinal injections on postnatal day (P)3–7. This age range was selected as only a small proportion of rods have degenerated(*18, 20*). We treated retinas with either RHO1-2A or B (2x10^10^ vg/eye), and assessed rod-isolated retinal function, after dark adaptation with full-field electroretinograms (ffERGs). RHO1-2A and B resurrected identical rod-isolated b-wave responses that were maintained through >140 days post injection (dpi; Fig. 3A and B) and, therefore, data from RHO1-2A and B (2x10^10^ vg/eye) are combined from Fig. 3C onward.

**Fig. 3.**
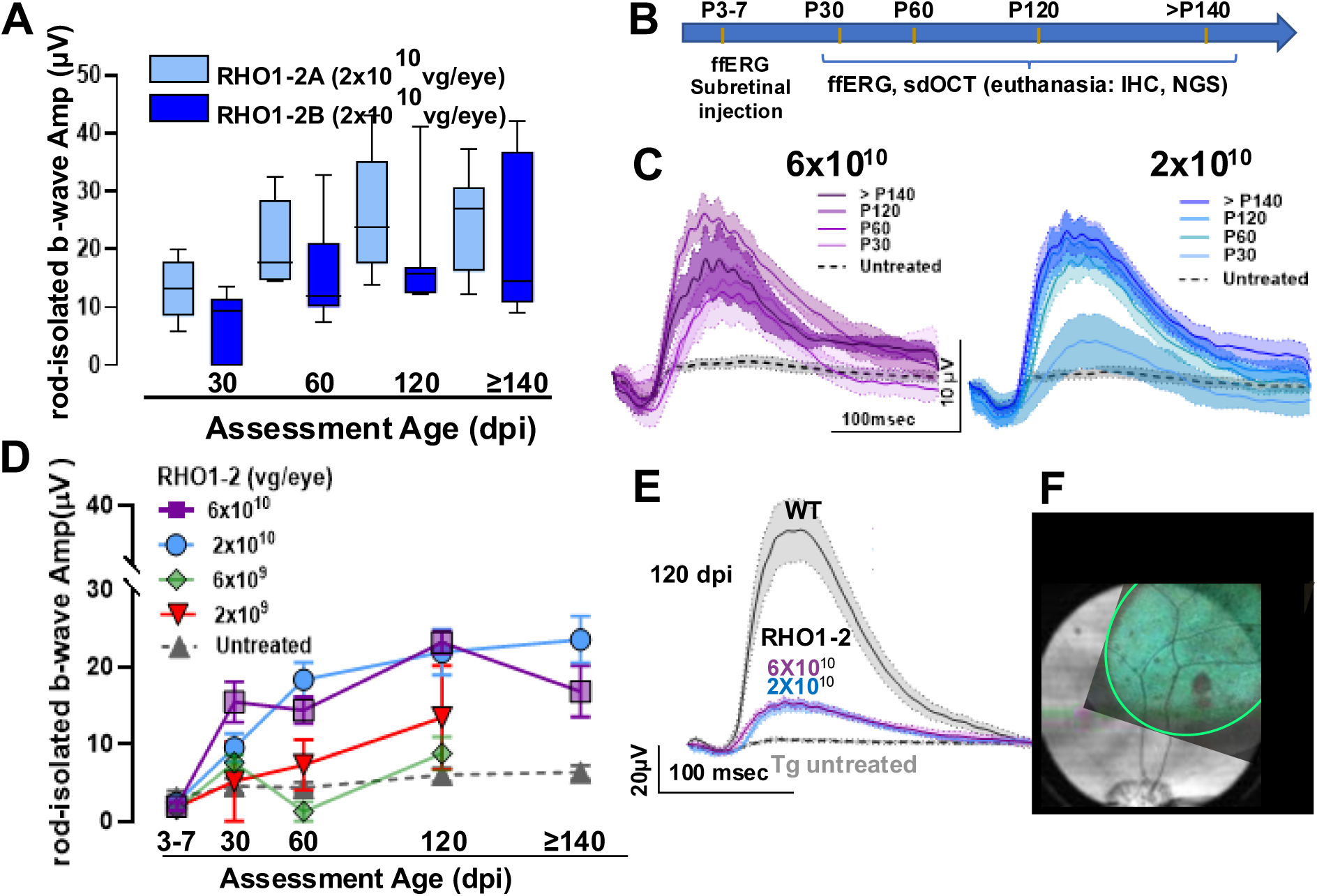
RHO1-2A and B treatment of TgP23H retinas is equally effective. (**A**) RHO1-2A and B (2x10^10^vg/eye) produce a similar resurrection of the rod-isolated b-wave in TgP23H pigs. (**B**) Dose escalation research design. Average rod-isolated ffERG b-waves at 6 and 2x10^10^vg/eye (**C**) across days post injection (dpi). Untreated age-matched Tgs have no response (grey dashed line, see Table 1 for numbers of eyes). (**D**) Dose escalation analysis of RHO1-2 treatment (P3-7) shows 6 and 2x10^10^ vg/eye produce significant rod-isolated b-waves compared to untreated from 30 to >140dpi (p< 0.03 to <0.0001; 2-way ANOVA, Tukey’s multiple comparison test). (**E**) Comparison of WT vs RHO1-2 treated (2 or 6x10^10^/eye) Tg rod-isolated b-waves at 120 dpi. (**F**) Superimposed fluorescence (green) and brightfield fundus images of a Tg retina (30dpi) injected with scAAV-GRK1:RHO1-2 and scAAV-GRK1:GFP illustrates the bleb size and location. Dark spot is the site of retinotomy.

**Table 1:** Table showing pigs treated under different conditions.

| RHO1-2 | P23H (+/-) | VirusTiter (vg/eye) | Treated eyes (n) | Untreated eyes (n) |
| --- | --- | --- | --- | --- |
| RHO1-2A | TG (N=6) | $2 \times 10^{10}$ | 6 | 4 |
| | | $1 \times 10^{10}$ | 2 | |
| RHO1-2B | TG (N=24) | $2 \times 10^{10}$ | 7 | 7 |
| | | $6 \times 10^{10}$ | 7 | 7 |
| | | $6 \times 10^9$ | 5 | 5 |
| | | $2 \times 10^9$ | 5 | 5 |

### RHO1-2 treatment resurrects a dose-dependent rod-isolated response in TgP23H pigs

We delivered one of four doses (2x10^9^ to 6x10^10^) of scAAV-GRK1-RHO1-2, spiked in scAAV-GRK1-GFP and delivered them subretinally to TgP23H pigs (P3-7, Table 1). Rod-isolated retinal function was resurrected in a dose-dependent manner. Responses were significantly larger in RHO1-2 treated Tg eyes at 2 or 6×10^10^ vg/eye compared to untreated age-matched Tg eyes. The two lower doses (2 or 6×10^9^ vg/eye) were similar to untreated Tg controls (Fig. 3C, D). At the minimum effective dose, 2×10^10^ vg/eye, the rescued rod response was significantly larger than untreated as early as 30dpi. We measured the extent of GFP expression in all RHO1-2 treated TgP23H pigs from fundus images. On average, the transduced area was 8x10mm, ∼10% of the retinal surface, which coincides with the percentage of the RHO1-2 treated Tg b-wave response relative to WT (∼10%; Fig. 3E, F). Therefore, RHO1-2 treatment resurrects a stable and dose-dependent rod-isolated b-wave response in Tg pigs that, without treatment, have no measurable rod-driven retinal activity (Fig. S2A, individual eyes).

### RHO1-2 treatment reduces outer nuclear layer (ONL) thinning and increases rod inner/outer segment lengths in TgP23H retina

We segmented retinal layers in sd-OCT b-scans and delineated RHO1-2^+^ and RHO1-2^-^ areas from aligned fluorescence fundus and sd-OCT brightfield images (Fig. 3F). A representative RHO1-2 treated TgP23H sd-OCT b-scan (Fig. 4A) shows a thicker outer retina (ONL+ inner + outer segments, IS/OS) at each RHO1-2**^+^** compared to each RHO1-2**^-^** location (Fig. 4A) or to untreated, age-matched Tg (similar retinal location; Fig. 4B). Within RHO1-2^+^ areas the outer retinal thickness was significantly thicker at each location in RHO1-2^+^ areas (1.4 to 5.6 mm from the transition zone) compared to RHO1-2**^-^** areas (same retinas; Fig. 4C, D) or to untreated age-matched Tg controls (similar locations; Fig. 4C) and remained same up to 300dpi (Fig. 4E, fig.S3A-C). The RHO1-2^+^ isolated ONL was significantly thicker (Fig. S3D, E) and inner retina layers were similar to WT.

**Fig. 4.**
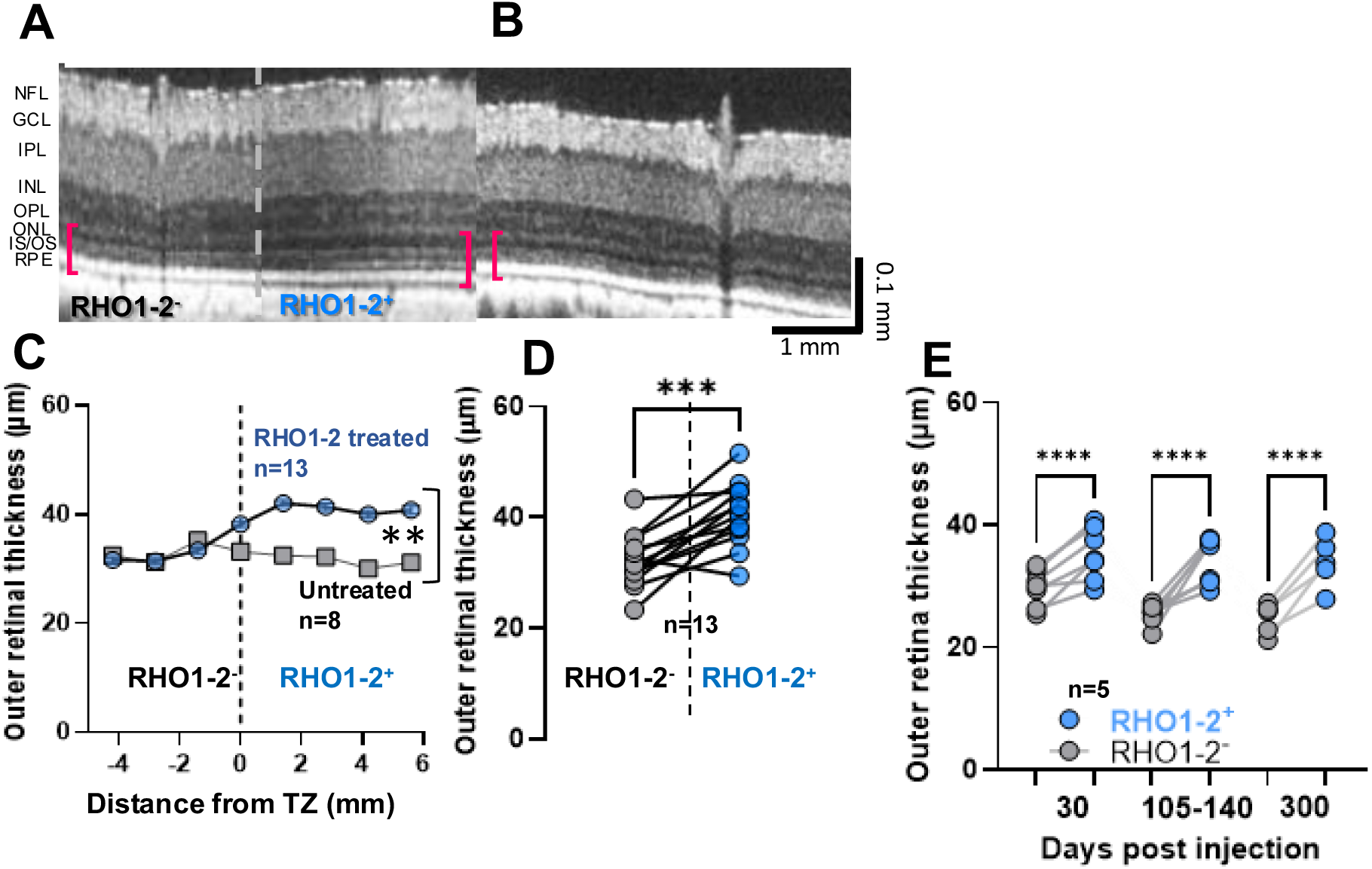
RHO1-2 treatment of TgP23H retinas reduces rod degeneration and increases outer segment length. (**A**) Representative sd-OCT b-scan of a TgP23H pig at 140dpi illustrates retinal layers within RHO1-2+ and RHO1-2-areas (right and left of the dashed line, respectively). Red brackets indicate the extent of the outer retina within RHO1-2+ and RHO1-2-areas. (**B**) Representative b-scan at a similar retinal location in an untreated age-matched Tg pig. (conventions as in A) (**C**) The outer retinal layers (ONL + OS/IS) within RHO1-2+ areas at 140 and 300dpi are similar (n=6 and 7, respectively) and are significantly thicker than untreated untreated Tg (C; p=0.009, unpaired t-test) or RHO1-2-areas (**D**; same retina; p=0.0002; paired t-test). (**E**) In five RHO1-2 treated Tg retinas (**Fig. S3 A-C**), average outer retinal thickness remains the same within RHO1-2+ or RHO1-2-areas. Thus, RHO1-2+ outer retina is significantly thicker across time (30 to 300dpi; p<0.0001; 2-way ANOVA, Sidak’s multiple comparison test).

Sd-OCT fundus images and corresponding b-scans were used to grade damage in RHO1-2 treated TgP23H retinas (Fig. S4). The site of the retinotomy was evident at 35dpi and did not change. Across eyes treated with RHO-2 ≤2x10^10^vg/eye (n=18), 89% showed by the retinotomy. In 11% of these eyes, we found damage limited to the outer retina surrounding the retinotomy (Fig. S4A and B). Among eyes treated with 6x10^10^vg/eye, 25% had damage throughout the retinal layers, of the rest, 50% had no damage and 25% confined damage (Fig. S4C).

### RHO1-2 treatment corrects rhodopsin mislocalization, elongates rod OS and reduces inflammatory marker expression in TgP23H pigs

We assessed RHO1-2 treated TgP23H rod morphology, using IHC and measured ONL thickness, rod morphology (GFP^+^) and rhodopsin expression (Fig. 5A and Table 3). Rods were rare in untreated and RHO1-2^-^ areas (Fig. 5B,C) in RHO1-2 treated retinas and were dysmorphic with mislocalized rhodopsin expression throughout the cell (Fig. 5C). At both 140 or 300dpi RHO1-2 treated retina had many rods with elongated IS/OS with rhodopsin expressed on the OS tips similar to WT (Fig. 5D vs. 5E). In RHO1-2 treated Tg retinas, the ONL was significantly thicker at each location in RHO1-2**^+^** areas compared to age-matched untreated retina (Fig. 5F) and to RHO1-2**^-^** areas (same retina; Fig. 5G).Within RHO1-2^+^ areas, IS/OS length significantly increased between 140 and 300dpi, where they approached WT length (Fig. 5H, dashed line).Thus, RHO1-2 treatment saves Tg rods from degeneration, corrects mislocalized rhodopsin expression, and leads to longer IS/OS, a result consistent with our functional data.

**Fig. 5.**
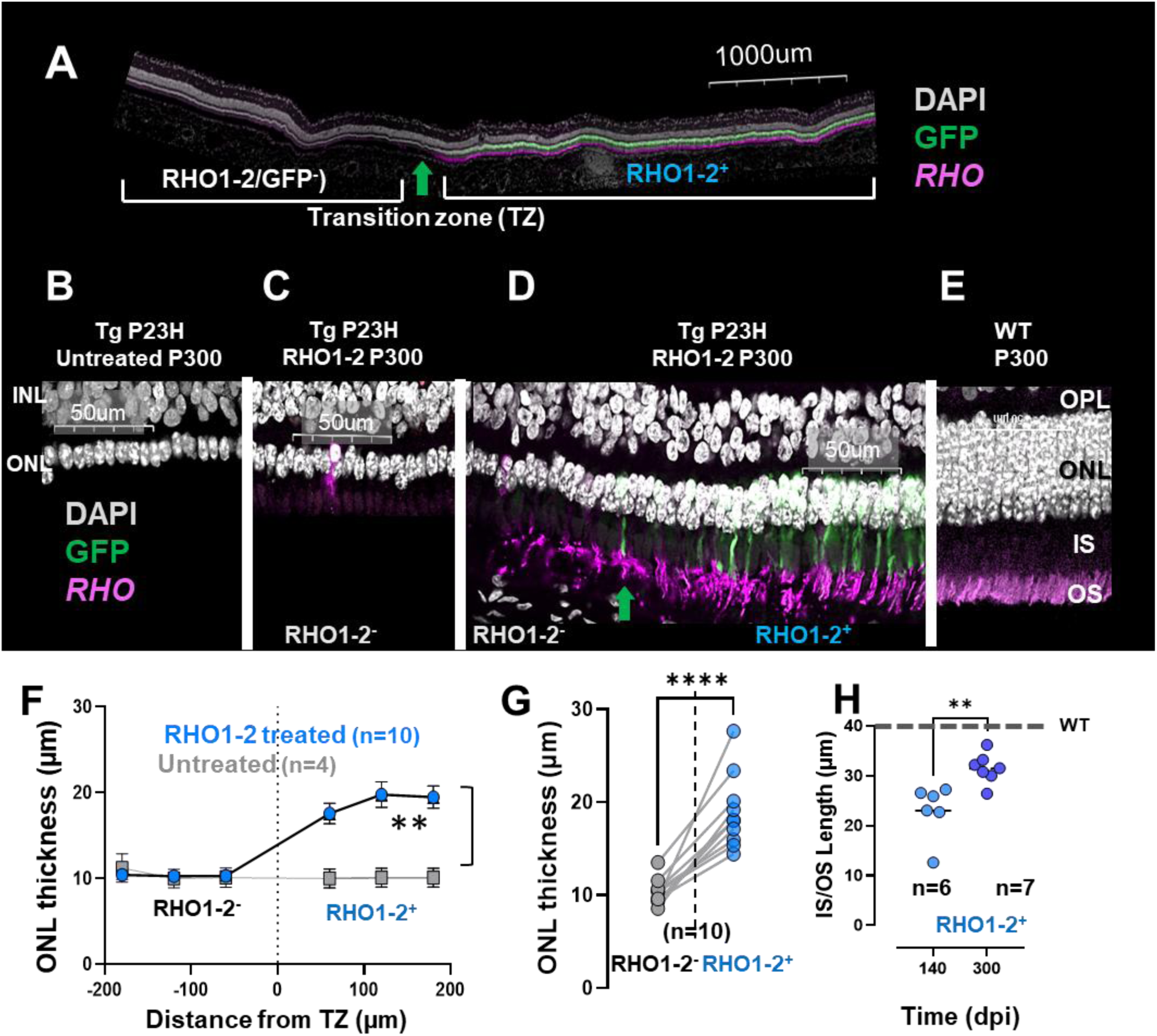
RHO1-2 treatment of TgP23H retinas retains ONL thickness, relocalizes rhodopsin, and regrows IS/OS. (**A**) Stitched confocal images of a RHO1-2 treated Tg retinal section (5mm in length) labeled for RHO (magenta), GFP(green), and DAPI (grey) illustrates the differences in ONL and rod morphology across RHO1-2+ and RHO1-2-areas (transition zone (TZ), green vertical arrow. Higher power confocal images from: an age-matched untreated Tg retina (**B**); RHO1-2-(**C**) and RHO1-2+ (**D**) areas in A and an age-matched WT pig (**E**). Untreated and RHO1-2-retinas have scattered RHO+ rods. In RHO1-2+ areas, RHO+ expression is limited to rod OS. (**F**) The ONL in RHO1-2+ areas is significantly thicker than untreated retinas (p=0.006; unpaired t-test) or RHO1-2-areas (same retina; **G**; p<0.0001; paired t-test). (**H**) Rod IS/OS in RHO1-2+ areas lengthen significantly with time (dpi) (p=0.005; Mann-Whitney test) and are almost as long as WT (dashed line).

RHO1-2^-^ areas and untreated Tg retinas showed a up-regulation of GFAP expression and migration of IBA1+ microglia into the ONL (Fig. S5A), like other adRP models(*22–24*). In contrast, within RHO1-2^+^ areas, GFAP and IBA1 expression were similar to WT (Fig. S5A, B), e.g., GFAP expression was localized to Mueller glia endfeet and IBA1^+^ microglia were excluded from the ONL.

### RHO1-2 treatment induces indels in the human mutant allele of TgP23H retinas

In cultured cells, RHO1-2 produced indels at the P23H target site (Fig. 2). To determine the nature of indels in RHO1-2 treated TgP23H pig retinas, we amplified a 231bp DNA fragment containing the c.68C>A site, sequenced and compared the PCR products from RHO1-2^+^ and RHO1-2^-^ areas. Results from a representative RHO1-2^+^ sample show that the highest fraction of indels occurred in RHO1-2^+^ samples at the RHO1-2 recognition sequence and frequency was dose-dependent (Fig. 6A - C). The most frequent indels were frameshift mutations (Fig. 6D) and non-coding indels were from the 5’ untranslated region of the amplicon. The predicted consequence of frameshift mutations is early termination of protein translation and in-frame indels are predicted to produce *RHO* with missense mutations. In both, the protein should not produce the P23H protein(*25*). The overall low fraction of reads with indels reflects the age of the pigs sampled (>250dpi), where the number of rescued rods in RHO1-2^+^ is low compared WT rods (Fig.5D vs E).

**Fig. 6.**
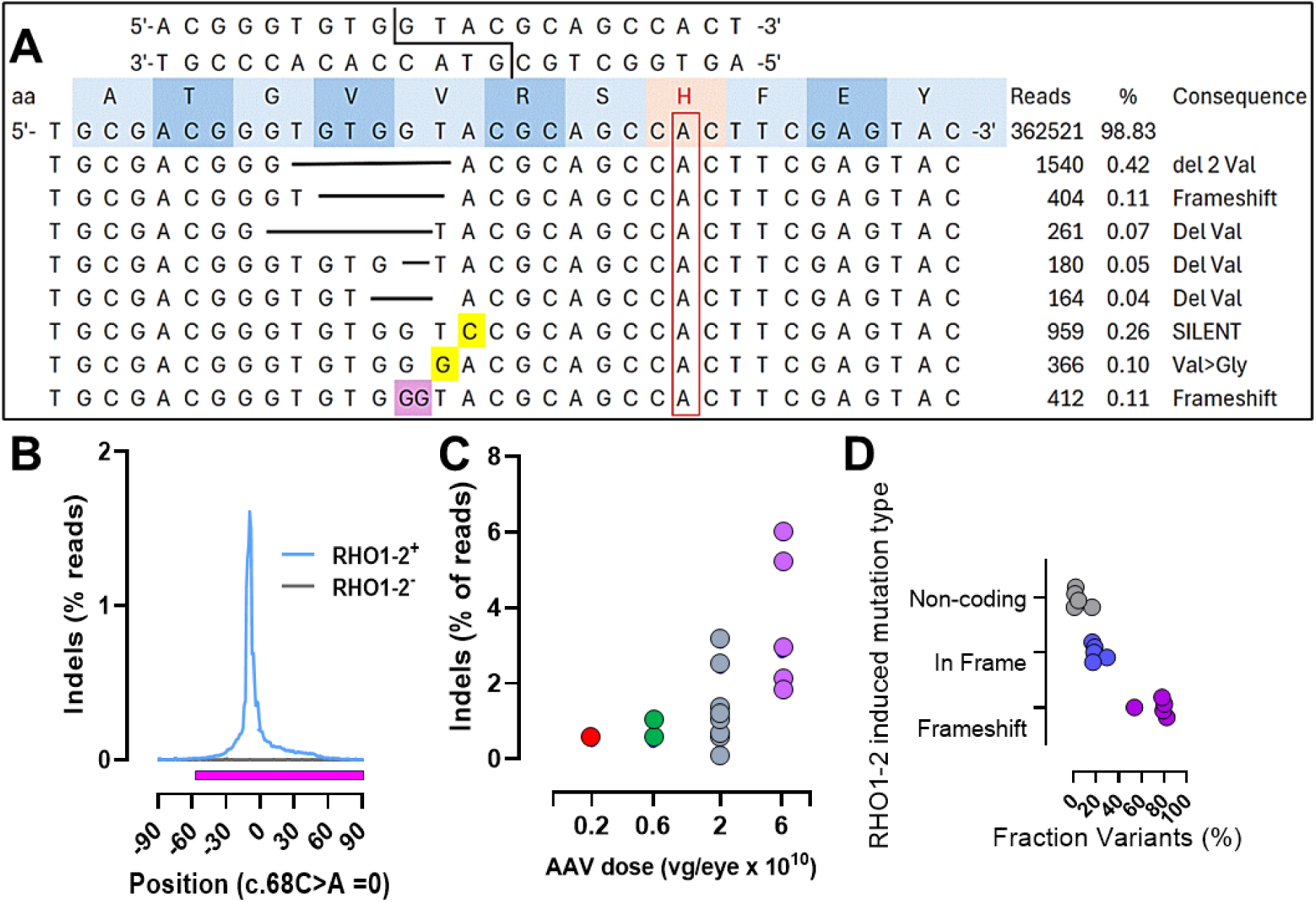
RHO1-2 treatment in vivo produces indels in the human P23H allele. (**A**) Sequence of the RHO1-2 recognition site showing cleavage and flanking sequence including the c.68C>A mutation (red box) producing the p.P23H protein. Top 8 most frequent mutations detected in AmpSeq data and their consequences. Horizontal bars indicate deletions. The purple shaded box indicates an inserted G, and the yellow boxes indicate base substitutions. (**B**) Indel frequency as a function of distance from the c,68C>A base in RHO1-2+ (blue) and adjacent RHO1-2-(grey) regions of a retina treated at P3 and analyzed at 60dpi. Magenta bar below the graph indicates the open reading frame. (**C**) Indel frequency increases in a dose-dependent manner (retinas >140dpi). (**D**) Mutation type for five samples from C. Non-coding are indels in the 5’ non-coding part of the amplicon (-123 to -1bp). In-frame and frameshift refer to indels in the ORF (0-109bp) within the amplicon.

### RHO1-2 treatment retains cone function and morphology in TgP23H pigs

In adRP, rod degeneration precedes cone dysfunction, leading to a current hypothesis that cone health depends on the presence of rods. We measured cone function using ffERGs with both a photopic flash and a 30Hz flicker stimulus (Fig. 7A, fig. S6A and B). Photopic b-wave amplitude was significantly greater in RHO1-2 treated compared to untreated Tg eyes at all times ≥60dpi, although the a-waves did not differ (Fig.7B). A significant difference in the flicker responses occurred at ≥140dpi (Fig. S6B).

**Fig. 7.**
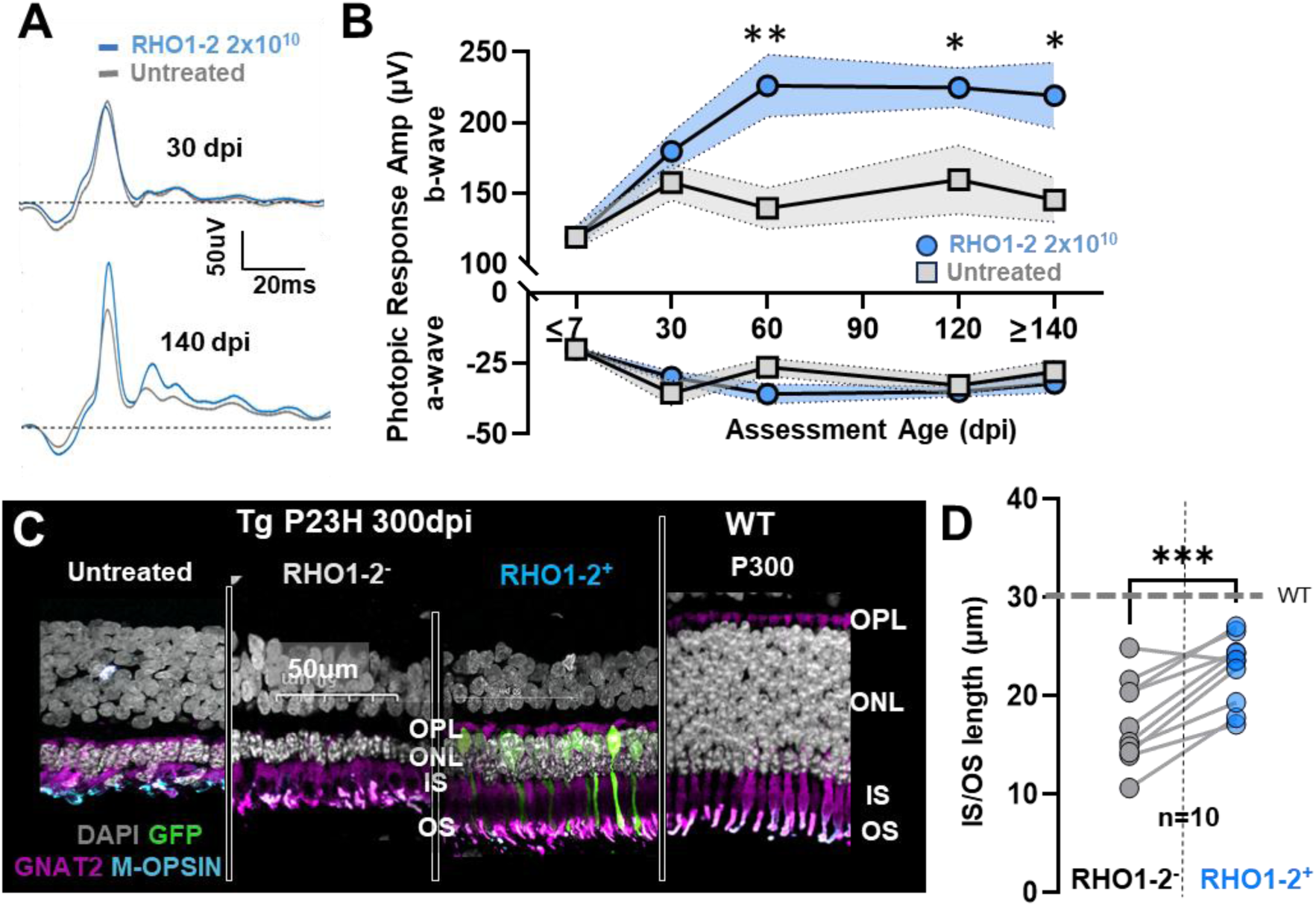
RHO1-2 treatment maintains cone function and morphology. (**A**) Representative cone dominated ffERG responses in RHO1-2 (2x10^10^ vg/eye treated and untreated TgP23H eyes. (**B**) Summary data showing that the a-wave is the same in RHO1-2 treated and untreated Tg eyes across time, and the b-wave is significantly larger in all RHO1-2 treated Tg from ≥ 60dpi (2-way ANOVA, Tukey’s multiple comparisons,**p=0.004;*p=0.04(120dpi);*0.02(≥140dpi). (**C**) Confocal images of age-matched untreated Tg retina, RHO1-2- and RHO1-2+ areas of RHO1-2 treated Tg retinas, and WT control. In RHO1-2-areas, Tg cones (GNAT+) IS/OS are stunted although m-opsin expression remains. In RHO1-2+ areas, cone morphology is maintained more similar to WT. Cone have long thin IS/OS similar to rods (GFP+) and display punctate m-opsin expression, similar to WT. (**D**) Cone IS/OS in RHO1-2+ areas are significantly longer than in RHO1-2-areas in the same retina (p=0.0003 ; paired t-test).

Cone morphology within RHO1-2**^+^** areas of TgP23H pigs was better preserved compared to RHO1-2^-^ areas, and similar locations in untreated Tg retinas (Fig. 7C). Inside RHO1-2**^+^** areas, cones maintained elongated IS/OS, with m-opsin expression on the OS tips, similar to WT (Fig. 7C). Untreated Tg cones (Fig. 7C) had swollen IS/OS, whose lengths were significantly stunted compared to those in RHO1-2^+^ areas or RHO1-2^-^ areas (Fig. S6D). Cone IS/OS in RHO1-2^+^ areas were significantly longer than in RHO1-2^-^ areas in the same retina (Fig.7D). Thus, cone function and structure are improved when RHO1-2 treatment rescues rods from degeneration and there may be a protective effect of RHO1-2 outside of the treatment area.

### RHO1-2 treatment enhances visual mobility under rod-isolated conditions in TgP23H pigs

A key metric of clinical efficacy is functional vision. We measured how quickly and accurately TgP23H pigs could use their RHO1-2 treated eyes to navigate a maze under photopic (20cd/m^2^) or rod-isolated (0.01 cd/m²) conditions (Fig. 8A). Several factors were used to ensure accuracy of our approach. Trials were spread over several days, the maze configuration was randomized to prevent memorization of the layout and experimenters were blinded to the experimental condition of each eye used to navigate the maze. We quantified several parameters that we hypothesized would be good indicators of vision, including speed (e.g., transit time, path length, steps to complete the maze) and accuracy (e.g., wrong turns and collisions with barriers). WT and TgP23H pigs were trained and tested binocularly under photopic conditions. One Tg pig had a RHO1-2 treated and an untreated eye, and its photopic performance was not assessed. The rest of the testing evaluated monocular performance by placing a light tight patch over one eye.

**Fig. 8.**
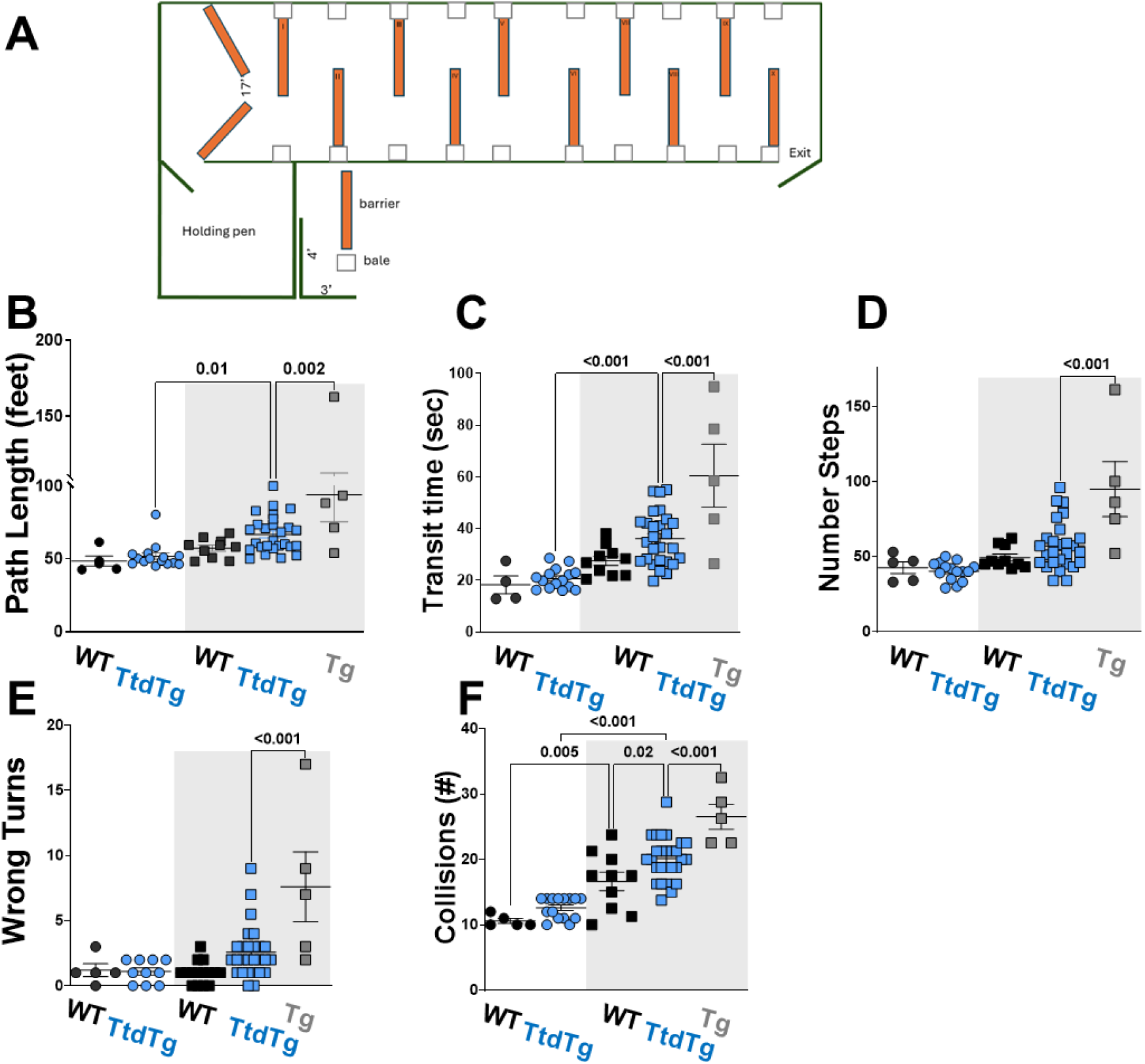
RHO1-2 treatment of TgP23H retinas improves rod-driven visual navigation. **(A)** Schematic diagram of one of three layouts of the maze used to test visual behavior. Orange plastic traffic barriers and hay bales (squares) could be moved to create three maze configurations. Behavior metrics recorded under photopic (white) and rod-isolated (grey) conditions that describe navigation speed and accuracy through the maze. Comparison of (**B**) Path length, (**C**) Transit Time, (**D**) Number of steps, (**E**) Wrong turns, and (**F**) Collisions of WT, RHO1-2 treated and untreated TgP23H eyes under photopic or rod-isolated (gray) conditions. Each point is a trial. WT (n=4 eyes), RHO1-2 treated (n=9 eyes), and untreated Tg eyes (n=1 eye). ANOVA with Sidak’s correction for multiple comparisons; all adjusted p values shown on graphs.

Under binocular photopic conditions, there were no differences in performance between WT and Rho1-2 treated TgP23H eyes (Fig. 8B-F, unshaded). WT pigs performance was similar under rod-isolated and photopic conditions on all parameters, although the number of collisions increased when only one eye was used to navigate. Under rod-isolated conditions (Fig.8B-F, grey shade), Tg pigs navigated the maze with their RHO1-2 treated eyes more slowly and less accurately compared to binocular photopic conditions. Here too there were more collisions with barriers. Despite the difference in performance between photopic and rod-isolated conditions, Tg pigs navigated the maze with their RHO1-2 treated eyes under rod-isolated conditions similar to WT. In the one Tg pig with a RHO1-2 treated and an untreated eye, navigation with the treated eye was significantly faster and more accurate than the untreated eye. These data show that RHO1-2 treatment of Tg eyes significantly improved functional vision in dim light.

## DISCUSSION

Retinitis pigmentosa (RP) is a group of IRDs that affect 1.5 million individuals worldwide(*26, 27*). Autosomal dominant RP (adRP) is a subset of these and is characterized initially by diminished night vision, followed by the loss of peripheral vision, and, ultimately, low vision of blindness. Currently there is no effective and approved treatment for this retinal degenerative disease available to the patients. While, the ultimate therapy will be mutation independent approaches to address the majority of RPs, these approaches remain challenging(*28*). Instead, an attractive approach for adRP would eliminate the mutant allele by cleavage, followed by host mediated DNA repair. This is especially amenable to P23H *RHO* adRP, for the most common form of adRP in North Americans (*29*), as a single WT *RHO* allele provides adequate visual function(*30*) and homozygous P23H *RHO* patients are so rare they are not described in the published literature.

Designer DNA endonucleases, exist in several formats, including ZFNs, TALENs, and CRISPR/Cas9, all of which demonstrate different degrees of specificity, promiscuity, and simplicity of design/engineering(*31, 32*). A feature common to all these endonucleases is that their components required for double stranded break induction are relatively large, making them difficult for gene delivery by AAV vectors with a packaging capacity of <5kb(*33*). The I-CreI meganuclease from *Chlamydomonas reinhardtii* overcomes the size limitations (920bp) and has high specificity because of this 22-24bp recognition sequence.

We show RHO1-2 meganuclease, re-engineered to specifically cleave the dominant negative P23H *RHO* allele is a potential therapy for adRP. Genome editing *in vitro* via RHO1-2A and B selectively eliminates the human P23H *RHO* mutant gene, leaving the human WT *RHO* gene intact. Indels were detected only at the on-target sites at the P23H target site and none were detected in the WT allele. Sequencing demonstrated that most edited mutant alleles have deletions >90%, of which >80% introduced an early stop codon, which would lead to premature translation termination and inactivation of the P23H *RHO* allele. Specificity analysis showed both RHO1-2A and B have very low genome wide off-targeting, 10 and 1 verified sites, respectively. For RHO1-2B, the one verified off-target site is located in an intergenic region more than 160 kb from coding genes and, therefore, would present a minimal risk of impacting any gene expression other than the intended P23H *RHO* allele.

We tested the efficacy and durability of RHO1-2 meganuclease genome editing in the only large animal model of human p.P23H *RHO* adRP (the transgene includes 4kb upstream of codon 1 to 7.5kb downstream of the stop codon of the human P23H *RHO* allele). From birth onward, these pigs have no rod-isolated retinal function, even though an almost complete complement of rods is present at birth. The absence of rod function provides a low background against which to evaluate early stages of genome editing. While the onset and time course of rod photoreceptor degeneration is more rapid than human disease, all of the human morphological and functional correlates can be found using sd-OCT imaging and ffERG recordings and analyses. We show RHO1-2 treatment of TgP23H pigs in early stage disease (P3 to 7) maintains ONL thickness, reduces rod degeneration and resurrects a rod-isolated ERG b-wave response, where none exists in untreated Tg pigs.

These Tg pigs carry multiple copies (8–12) of the transgene and we do not know how many are functional. We are currently using PacBio and ONT LongRead Sequencing to compile the entire transgene inserted on Chromosome 5. With this in mind and because the pig eye is similar in size to human, our results indicating that 2X10^10^ vg/eye, the minimal effective dose in the pig, provides an upper estimate of a clinically relevant dose for patients that only have a single mutant copy. This means a RHO1-2 therapeutic approach will be even safer in patients in the clinic. In addition to functional efficacy, our sd-OCT (*in vivo*) and IHC (*ex vivo*) show morphological rescue of rods themselves. We are currently developing AI/machine learning to correlate outer retinal sd-OCT banding patterns with cellular morphology to better detect what is maintained with RHO1-2 treatment in clinical applications with patients.

The relocalization of RHO expression as well as the elongation of IS/OS in RHO1-2 treated retinas suggest that elimination of P23H *RHO* expression allows WT *RHO* (pig *RHO*) to traffic properly and rebuild stunted IS/OS. While aspects of the molecular mechanisms that cause P23H *RHO* adRP remain under study, our results support the idea that misfolded P23H opsin disrupts disc formation/organization of rod outer segments(*34, 35*), although we cannot rule out a role for misfolded rhodopsin as a trigger of rod death.

RHO1-2 rescue of rods maintains cone function, morphology (IS/OS lengths), and opsin expression. Together with the significant difference in the IS/OS length of cones in adjacent RHO1-2^-^ areas vs untreated cones supports the hypothesis that a diffusible factor, e.g., RDCVF, is required for cone health(*36*).

RHO1-2 mediated improvements in retinal structure and function persisted without decline for at least a year post-treatment. This efficacy extends to visual behavior. WT and TgP23H pigs used their RHO1-2 treated eyes to navigate through a maze under rod-isolated conditions with nearly equal speed and accuracy. Using their treated eyes, Tg pigs navigate the maze better under rod-isolated conditions than with an untreated eye. Both WT and RHO1-2 treated Tg pigs collide with barriers significantly more frequently when they navigate with only one eye. RHO1-2 treated Tg pigs’ accuracy also may be limited by the small area of rescued retina (∼10% of total). Thus, limiting the visual field leads to more collisions. The difference between RHO1-2 treated and untreated Tg eyes also may indicate that the presence of central rods may have a therapeutic influence on peripheral rods.

We evaluated the specificity of the RHO1-2 at the pig P23H targeting site, but because the pigs do not have a human WT allele, we cannot address RHO1-2 cleavage of the WT pig allele *in vivo*. We also did not examine off-target cleavage of RHO1-2 in pigs, because the DNA architecture is unlikely to be the same as humans. A human retinal organoid system will be used in the future.

Although our data suggest that RHO1-2 treatment for P23H adRP should be therapeutic, a greater understanding of its potential risks and benefits is still needed before proceeding to clinical trials. Further studies, may be needed in non-human primates to better evaluate potential inflammatory reactions, even though the pig immune system is relatively similar to human(*37*). New delivery vectors or non-viral delivery may be needed for optimized delivery in patients. Ex vivo retinal cultures, human WT or P23H *RHO* retinal organoids may be useful in studying the activity of RHO1-2B and further verifying cleavage at off-target sites in a setting even more similar to our *in vivo* results.

Taken together, our results provide evidence that engineered meganucleases can be tailored to target the *RHO* P23H allele, creating a strong basis for exploring new strategies for human dominant diseases and eventually successful clinical application and treatment/cure for patients with P23H adRP.

## MATERIALS AND METHODS

### Nuclease generation

Methods for generating target specific nucleases have been described previously(*38*). Briefly, I-CreI libraries used in directed evolution, two I-CreI monomers were generated that were predicted to recognize each half of the P23H *RHO* target sequence (5’-<u>ACGGGTGTG</u>GTAC<u>GCAGCC**A**CT</u>, half-sites underlined, mutant base bolded) and fused onto a single peptide sequence library. The fused library of RHO1 and RHO2 monomers was selected in directed evolution to cut the P23H *RHO* target sequence. Further, they were counter selected so that they did not cut the WT *RHO* sequence, or other closely related sequences. The resulting nucleases were screened in a CHO GFP assay for activity against the P23H *RHO* target and for discrimination against the WT *RHO* sequence. The best initial candidate, RHO1-2A, was evaluated for off-target cutting of the WT allele sequence. Amino acids predicted to be site discriminating P23H versus potential off-targets including the WT alleles were subsequently rerandomized and used in further directed evolution procedures to produce RHO1-2B. All plasmids used in nuclease generation were custom-made.

### Creation and characterization of CHO reporter cells

As described(*38*), CHO-K1 Flp-InTM cells (ThermoFisher Scientific; R75807) were produced and maintained according to the manufacturer’s protocol and used to create our meganuclease reporting cell lines. Two plasmids were created containing our positive control recognition sequence (Meg-23/24 meganuclease) and either the P23H mutant recognition sequence or the WT sequence (Fig 2 Ai, Aii, respectively). The reporter cassette comprised, in 5’ to 3’ order: an SV40 Early Promoter; the 5’ 2/3 of the GFP gene; the recognition sequence (P23H or WT RHO); Meg-23/24 meganuclease; and the 3’ 2/3 of the GFP gene. CHO-K1-Flp-In TM cells were co-transfected with one of these two reporter cassette plasmids and the Flp recombinase. Following transfection and a short recovery, they were selected via antibiotic resistance for correct insertion of the cassette into the genome at the Flp / FRT site. Cleavage of any of the target sites results in single-strand annealing (SSA) repair, reconstituting a functional GFP gene.

### Evaluation of RHO1-2 meganuclease editing of CHO reporter cells

To assay the specificity of RHO1-2 meganucleases, 5x10^4^ CHO reporter cells were transfected with 90 ng of RHO1-2A, RHO1-2B, HPP1-2 (an unrelated ARCUS nuclease that targets a site in plants and is used as a mock), or Meg-23/24 meganuclease (positive control) mRNA in a 96-well plate using Lipofectamine® MessengerMax (ThermoFisher) according to the manufacturer’s instructions. The number of cells expressing GFP, an indicator of cleavage, was measured by flow cytometry. Transfected CHO reporter cells were cultured for 24 hours, then dissociated and re-suspended in 3% fetal bovine serum (FBS in PBS), split for flow on day 2, and then similarly dissociated and re-suspended for flow on day 5 post-transfection. The percentage of GFP-positive CHO cells was measured and compared to an untransfected negative control at each time and was normalized to the % GFP-positive cells produced by Meg-23/24 meganuclease.

### Generation of c.68C>A mutation (P23H) in human K562 cells

We created a human cell line carrying both WT and P23H *RHO* alleles using chronic myelogenous leukemia cells (K562; ECACC; <u>89121407</u>). We cotransfected (Lonza Nucleofector 2B) 9.17x10^5^ K562 cells/ml with 3.5µg of ARCA capped RHO3-4 mRNA, encoding a nuclease engineered to recognize the WT *RHO* sequence at the equivalent location as RHO1-2 and a 160nt P23H *RHO* donor single-stranded template (Integrated DNA Technologies, IDT). Rho3-4 induced a double-strand break at position 68, and DNA repair used the 160nt sequence as a template replacing the ‘C’ with an ‘A’ at position the c.68C>A, creating the P23H *RHO* allele. To identify colonies bearing the c.68 C>A transversion (P23H RHO), cells were cultured and dissociated 48 hours after co-transfection and serially diluted to a final concentration of 0.3 cells/100 μL. Approximately 100μL of diluted cells were plated into each well of nine 96-well plates, incubated in 5% CO_2_ at 37°C until confluent, and their genomic DNA extracted (Quick Extract DNA kit). 192 clones were identified by digital droplet PCR (ddPCR) containing the P23H *RHO* mutation and were validated by Sanger sequencing. One clone, named 2G11, carrying the WT and P23H alleles, was expanded and maintained in RPMI 1640 (Thermofisher; 11875135) supplemented with 10% FBS and in a 5% CO_2_ atmosphere at 37 °C.

### Evaluation of RHO1-2 editing of P23H and discrimination of WT alleles in 2G11 cells

To evaluate RHO1-2 nuclease specificity for the P23H *RHO* compared to the WT *RHO* allelle, 1x10^6^ 2G11 cells were electroporated with 5μg mRNA encoding the RHO1-2 nuclease (Lonza Nucleofector 2B), cultured for 2 or 5 days after electroporation, and pelleted. The genomic DNA was isolated for digital droplet PCR (ddPCR) analysis (Macherey Nagel NucleoSpin Blood Quickpure Kit).

### Digital droplet PCR

ddPCR was performed using a QX200 ddPCR system (Bio-Rad) according to the manufacturer’s instructions. Briefly, 120ng of purified genomic DNA was combined with 12µl ddPCR Supermix for Probes (no dUTP; Bio-Rad), 1.2µl of the 20X master mix for the amplicon (final concentration in the reaction: 900nM of each primer, 250nM of a fluorophore-conjugated probe), and 0.25µl 20 U/ml of *HindIII-HF* restriction endonuclease (New England Biolabs) in nuclease-free water to a final reaction volume of 24μL. Samples were partitioned into approximately 20,000nL-sized droplets (QX200 droplet generator (Bio-Rad) and PCR performed on a C1000 Touch thermal cycler (Bio-Rad) using a three-step cycling profile: 10min at 95°C, 45 X (30s at 94°C, 30s at 60°C, 2min at 72°C), and 10 min at 98°C. Once the cycling was completed, acquisition was performed using absolute counts, and the probes were assigned to the correct dye (FAM for P23H *RHO* and VIC for WT RHO). Droplets were gated, positive and negative droplets were distinguished in both channels (QuantaSoft Analysis Pro (BioRad)), and the ratio of P23H to WT *RHO* positive droplets was calculated.

### scAAV plasmid, virus production, and characterization

Self-complementary AAV plasmids encoding the RHO1-2 nuclease or GFP included either the ubiquitous cytomegalovirus (CMV) or the photoreceptor-restricted G-protein coupled receptor kinase 1 (GRK1) promoter(*39*). Plasmids were purified (Macherey Nagel Maxi Plasmid Preparation Kit), and self-complementary recombinant AAV5 (scAAV5) was produced in HEK293 cells using a triple transfection protocol, followed by CsCl gradient centrifugation and dialysis in 1× phosphate-buffered saline (PBS) solution as previously described. For each viral preparation, physical titers (viral genome (vg/ml) were determined by quantitative real-time PCR (Roche LightCycler 480 real-time PCR) and further confirmed by comparison of the Southern dot blot of DNA prepared from purified viral stocks and defined plasmid controls as described previously(*38*).

### Virus preparation

The virus was prepared in a sterile hood on the day of the subretinal injections. scAAV5-GRK1-RHO1-2A or RHO1-2B was spiked with scAAV5-GRK1-GFP (3:1 ratio) and diluted in 0.001% Pluronic F68 Dulbecco’s phosphate-buffered saline (DPBS) to the appropriate viral titer (Table 1). The virus was loaded into a microdose injection kit (MedOne Surgical, Inc., Sarasota, FL, USA) attached to a viscous fluid control pack (Alcon, Inc., Fort Worth, TX, USA) connected to a vitrectomy machine (Constellation system, Alcon Inc, Fort Worth, TX, USA) for precise injection of the virus volume.

### Targeted deep DNA sequencing

Amplicons, generated using Phusion II Hot Start Polymerase (ThermoFisher), with forward primer i5-CTTCGCAGCATTCTTGGGTG, reverse primer CCCATCAACTTCCTCACGCT-i7, and were sequenced on an Illumina MiSeq (2x150bp paired end run) sequencer. Reads 1 and 2 were merged using Flash(*40*) and aligned to the reference sequence using BWA-MEM(*41*). Insertions and deletions were identified using the CIGAR (Concise Idiosyncratic Gapped Alignment Report) and characterized as in or out of frame changes using either publicly available(*42*) or custom software.

### Oligo Capture off-target site identification

2G11 cells (1x10^6^) were electroporated with a double-stranded oligo containing randomized four bp 3’ overhangs on both ends (500ng) and either RHO1-2A or RHO1-2B mRNA (1µg). After 2 days in culture, cells were pelleted, and genomic DNA (gDNA) was isolated (Macherey Nagel NucleoSpin Blood Quickpure kit). The gDNA was fragmented (Covaris focused ultrasonicator) and ligated to an adapter containing unique molecular identifiers (UMI). A two-step nested PCR strategy amplified the DNA between any inserted oligo and ligated adapter, which was sequenced (Illumina NextSeq 2000; (2x150bp paired end run)). Reads were aligned to the reference genome (GRCh38) and PCR duplicates removed using the aligned positions and UMIs. Oligo insertion sites were determined, as was the most likely nuclease cut site within 50bp of the insertion site, using custom software. Oligo capture was repeated three times on independent biological replicates for RHO1-2A and RHO1-2B nucleases to identify off-target sites.

### rhAmpSeq verification of potential off-targets

A rhAmpSeq panel containing primers for the on-target site and the 38 putative off-target sites identified in OligoCapture was ordered (IDT/DNA). 2G11 cells (1x10^6^) were electroporated with either 500ng or 1000ng RHO1-2A or RHO1-2B mRNA. After 2 days in culture, cells were pelleted and gDNA isolated. rhAmpSeq amplification was performed using 200ng of gDNA per sample (IDT protocol; rhAmpSeq library preparation for Targeted Amplicon Sequencing). The resulting libraries were sequenced on an Illumina NextSeq2000 (2x150bp paired-end run) and analyzed for the presence of indels within the 22bp RHO1-2 cut site using custom software.

### RHO1-2 antibody production

A rabbit polyclonal antibody was produced against a purified I-CreI RHO1-2.

### Animals

All protocols were approved by the University of Louisville Institutional Animal Care and Use Committee and adhere to the ARVO Statement for Use of Animals in Ophthalmic and Vision Research. Piglets were born and housed in the University of Louisville AALAC certified facility with controlled 14/10 hour light/dark cycle. After weaning, pigs were fed twice daily and provided water ad libitum. Pigs were socially housed and cage size and number housed/cage dictated by USDA guidelines. Daily observations of the pigs’ health were made. Enrichment was provided and changed regularly. TgP23H and WT piglets were created by artificially inseminating wild type (WT) domestic sows with semen from a TgP23H miniswine boar (rrid-NSRRC:0017). At ∼postnatal day 2 (P2; P0 is the day of birth), a small tissue biopsy was used to isolate DNA for genotyping(*43*). Genotypes were verified by the absence/presence of a rod-isolated full-field electroretinogram (ffERG; described below at P3-7).

### Animal preparation for aseptic subretinal surgery, ffERG, and OCT assessments

Animal preparation for full-field ERGs has been published(*20*). Piglets <P14 were sedated and anesthetized via intramuscular (IM)/subcutaneous (SC) administration of a mixture of Ketamine (5mg/kg), Butorphanol (0.2mg/kg), Dexmedetomidine (0.002mg/kg), and Midazolam (0.2 mg/kg). Piglets >P14 were sedated and anesthetized with a mixture of Ketamine (5mg/kg), Dexmedetomidine (0.02mg/kg), and Midazolam (0.2 mg/kg) via IM/SC route. In all pigs, anesthesia was maintained with 1-3% isofluorane delivered via an endotracheal tube. Vital signs (e.g., SpO2, respiratory rate, heart rate, blood pressure, and body temperature) were monitored every 5 mins throughout the procedure. An IV catheter, placed in the ear vein, delivered intravenous fluids (Lactated Ringers Solution with or without 5% dextrose; 10-15ml/kg/hr) to maintain blood pressure and normal glycemic levels (60–140 mg/dL). Anesthetic levels were adjusted to sustain respiration and heart rate within normal levels. Pupils were dilated and accommodation relaxed with topical application of 2.5% Phenylephrine Hydrochloride (Paragon Bioteck Inc., Bridgewater, NJ, USA) and 1% Tropicamide drops (Bausch and Lomb, Bridgewater, NJ, USA). Adjustable lid specula kept the eyelids separated during testing. Saline drops were administered as needed to maintain corneal lubrication and clarity.

### Subretinal injections of AAV-RHO1-2A and 2B

One to two days before subretinal surgery, piglets were orally dosed with flavored prednisone (dose: 0.5-1mg/kg, Wedgewood Pharmacy LLC, Lot #000-06831886), which was continued for 30 days, with a one-week tapering dose administered every other day. A sterile vitreoretinal surgical approach was used to access the subretinal space (between the retina and the pigment epithelium) and two 25g trocars (Alcon, Inc., Fort Worth, TX, USA) were placed at 2-2.5mm posterior to the limbus, one in the superior-nasal and the other in the inferior-nasal quadrant. A light pipe inserted into one trocar visualized the retina and a 25g subretinal cannula with a 41g flexible tip was placed into the other trocar. A local retinal detachment (bleb) was created, ∼50 μL total volume of virus or saline (control) was injected into the subretinal space, which created an average bleb 8x10mm. After the injection, the needle and trocars were removed, trocar sites were dabbed with sterile cotton applicators to promote self-healing and antibiotic and steroid ointments placed topically.

### Clinical examination and fundus imaging

Clinical examinations were performed pre- and at regular intervals post-injection and included slit lamp biomicroscopy (Kowa company Ltd., Torrance, CA, USA) to assess the anterior segment and indirect ophthalmoscopy (Keeler, Malvern, PA, USA) to assess the fundus. Restraint was used in piglets (<P30), and thereafter, sedated with Ketamine (5mg/kg), Dexmedetomidine (0.02mg/kg), and Midazolam (0.2mg/kg). Eyes were evaluated for: cataract formation, retinal tears, inflammation, optic nerve pallor, retinal arterial vasculature, and migration of pigment (bone spicules). Following clinical examination, fundus fluorescence images were collected (Micron X imaging system; Phoenix Research Labs, Pleasanton, CA, USA) to determine the onset and extent of GFP expression. We also evaluated the effect of subretinal injection on the area surrounding the retinotomy (Fig.S4).

### ffERG recording and analysis

ffERG methods have been described previously(*20*), Flash stimuli were produced, and responses recorded using a UTAS ERG system with a BigShot Ganzfeld stimulator (LKC Technologies, Inc., Gaithersburg, MD, USA.). The head was placed inside the Ganzfeld bowl, or when the pig’s heads were too large for the Ganzfeld, a hand-held ERG system (Retevet; LKC Technologies, Inc., Gaithersburg, MD, USA) was used. Bilateral ERGs were recorded using JET electrodes (The Electrode Store, Enumclaw, WA, USA) that were wetted with 2.5% hypromellose solution (Hub Pharmaceuticals LLC, Farmington Hills, MI, USA) and placed on the cornea. A ground electrode was placed on the midline of the forehead, and reference electrodes were placed in the soft tissue above the orbit of each eye.

Table 2 shows the details of the scotopic and photopic flash stimulus protocol, modified from the ISCEV standard for clinical ffERG. Rod-isolated and scotopic responses were collected after 20 minutes of dark adaptation. Photopic and cone isolated (30 Hz flicker) responses were collected after 5 min of light adaptation (20cd·m^-2^ background). Individual responses to each stimulus within a trial were analyzed using a custom MATLAB script, aberrant waveforms were rejected, and the remaining responses averaged. The averaged a-wave amplitude was measured from baseline to trough within a 40msec window beginning at stimulus onset. The rod-isolated response has no a-wave and b-wave amplitude was measured from baseline to peak within a 90msec window beginning 60msec after stimulus onset. The photopic response was measured from the a-wave trough to peak using the same timeframe.

**Table 2:** Custom ISCEV Electroretinogram (ERG) protocol.

| DARK ADAPTATION : 20 MINUTES |  |  |  |  |  |
| --- | --- | --- | --- | --- | --- |
| Step | Test Sessions | Flash intensity (cd.s/m<sup>2</sup>) | Background (cd/m<sup>2</sup>) | Number of flashes | Time interval between flashes (sec) |
| 1 | Scotopic -Rods only | 0.01 | none | 15 | 0.5 |
| LIGHT ADAPTATION : 5 MINUTES; BACKGROUND: 20 cd.s/m<sup>2</sup> |  |  |  |  |  |
| 2 | Standard Cones | 3 | 20 | 30 | 0.5 |
| 3 | Flicker | 3 | none | 210 | 0.033 |

### Spectral domain OCT (sd-OCT) Imaging and analysis

sd-OCTs were acquired in anesthetized pigs at ≥P21 (breathing artifacts in young pigs prohibit collection of high-quality images) using a Bioptigen Envisu R2200 SD-OCT (Leica Microsystems Inc., Buffalo Grove, IL) with a broadband light source (central wavelength 878.4 nm, 186.3 nm bandwidth; Superlum, Enterprise Park, Cork, Ireland). Eye position was stabilized and manipulated, using scleral stay sutures, placed on the conjunctiva oriented parallel to and just posterior to the limbus along the superior-inferior and nasal-temporal axes. Images were acquired using the InvivoVue software (Bioptigen). The reference arm position was calibrated and fixed throughout the scans to procure high-quality images at appropriate scanning depths. A focus correction adjustment (range of +8 to −20 D) enhanced scan sharpness. A 14 mm X 14 mm rectangular volume was imaged, centered on the visual streak and encompassing the injection site and GFP^+^ areas inside and GFP^-^ areas. Frames from sd-OCT b-scans (1000 A-scans/B-scan, 10 B-scans averaged sequentially, giving 100 B-scans) were acquired under optimized illumination settings (Gain value of -1 to +1). Acquired sd-OCT images were averaged offline (InVivoVue software) to detect the hyper- and hypo-reflective bands of the outer retina (using calipers), and quantify ONL and IS/OS thickness, from the distance between the border of the hyper-reflective OPL to the hyper-reflective RPE. We took four OCT b-scans through the bleb area (at an interval of ∼1.5-2mm) between the superior and inferior extent of the RHO1-2^+^ area (∼8mmx 8mm), which also contained a transition zone to RHO1-2^-^ areas on the fundus image) or similar areas in untreated retina. We measured the outer retinal thickness across the entire b-scan at 1.4 mm intervals using Diver (InVivoVue software). Measurement points were placed at retinal layer transition zones to delineate the borders of the ONL, external limiting membrane, IS/OS and RPE(*44*). We plotted the outer retinal width as a function of distance from the transition zone. To generate heatmaps (Fig.S4) depicting retinal layer thickness, we converted b-scan layer segmentations into 2D representations of volume-wise layer thickness. First, the entire set of averaged b-scans, their layer segmentations were saved in a single OME TIFF volume and converted into thickness values by finding the single pixel wide column wise midline of each layer. In a new empty volume, the pixel value at each of the midline locations was changed to the layer thickness in micrometers of that column. This process was repeated for every b-scan and every layer in the full set of 100 average b-scans. Finally, the per layer XZ maximum intensity projections of this thickness volume were created and saved to OME-TIFFs, creating the final heatmaps of layer thickness where the pixel brightness value is the layer thickness in micrometers.

### Tissue processing and Immunohistochemistry (IHC)

Methods for tissue processing have also been previously published(*20*). At each terminal assessment point, a surgical plane of anesthesia was induced, and the eyes were enucleated to maintain high-quality RNA. Immediately after, the pig was euthanized with an intravenous injection of a pentobarbital-based solution (Euthanasol, cat # 004988, Virbac Corp, Westlake, TX). Vital signs were monitored until respiration and cardiac function ceased, and a thoracotomy was performed. The posterior pole of the eye was isolated, and GFP fluorescence was imaged across the retina using a fluorescence, stereo zoom microscope (Olympus SZX16 Stereo high resolution microscope). The retina was removed, flattened, and pieces dissected based on retinal location.

For IHC, retinal pieces were dissected to contain RHO1-2^+^ and RHO1-2^-^ areas for comparisons across infected and uninfected retina. Pieces were fixed in 4% paraformaldehyde in 0.1 M phosphate buffer (PBS) for 20 minutes at room temperature (RT), followed by three washes with 0.1 M PBS. The tissue was cryoprotected through a sucrose gradient of 5%, 10%, 15% (in PBS; 1 hour each, RT), and 20% sucrose overnight (4 °C). Retinas were incubated in an OCT/20% Sucrose (2/1) solution (RT), placed in molds containing the same solution, and frozen over a liquid nitrogen-cooled 2-methyl butane bath. In most cases, two to three retinas were embedded in the same mold so that infected/control areas or pigs in the same or different conditions could be reacted and compared on the same slide. Frozen sections were cut (20μm) on a cryostat (CM1850, Leica Biosystems Inc., Buffalo Grove, IL), placed on Super-Frost glass slides (Fisher Scientific, Pittsburgh, PA), and stored at -80 °C. Before IHC reactions, frozen sections were thawed and dried at 37°C for ∼60 minutes and rinsed in PBS for 5 mins. For permeabilization, sections were incubated in 0.5% Triton X-100 in PB (PBX; 15 mins, RT), and blocked with 5% donkey serum in PBS (1 hour, RT). Following removal of the blocking solution, sections were incubated with primary antibodies diluted in blocking solution (overnight, 4°C). Sections were washed three times with PBS (10 mins/wash) and reacted with secondary antibodies diluted in PBS (1 hr, RT). Table 3 provides the details of the primary and secondary antibodies. Sections were imaged on an Olympus FV3000 or 4000 confocal microscope (Olympus Confocal America, Inc., Center Valley, PA), using a 10x (NA=0.45) and 40x water/oil objective (NA=1.45). The images shown are full projections of z-stacks 5 – 6μm total depth, which was chosen as 5μm is the diameter of photoreceptor nuclei, and this thickness provides images without significant overlap of adjacent nuclei/IS/OS. All sections were imaged with the same confocal settings, and if adjustments to contrast and brightness were needed, they were made using the same settings for all sections (FluoView software; FV31S-SW, Olympus Confocal America, Inc., Center Valley, PA). From these images, we measured the ONL thickness (from OPL to photoreceptor IS border) and IS/OS length using Olympus CellSens Software (Olympus Confocal America, Inc., Center Valley, PA). We measured thickness at six different positions for each retinal section, spaced 60 µm apart. The thickness of areas with and without GFP expression was separated, and the average thickness was plotted as a function of the distance from the transition zone of RHO1-2^+^ and RHO1-2^-^ areas.

**Table 3:** Primary and Secondary Antibodies used in Immunohistochemistry (in Pigs)

| Antibody | Host Species | Working dilution | Cat# | Manufacturer/Gift of |
| --- | --- | --- | --- | --- |
| **Primary** |  |  |  |  |
| Rhodopsin-clone1D4 | Mouse | 1:1000 | # **MA1722** | Thermofisher Scientific, Waltham, MA, USA |
| REEP6 Polyclonal | Rabbit | 1:500 | #**12088-1-AP** | Proteintech Group, Inc, Rosemont, IL, USA |
| S-opsin | Chicken | 1:500 | # **JH008** | Kerafast Inc, Boston, MA, USA |
| L/M-opsin | Rabbit | 1:500 | # **AB5405** | MilliporeSigma, Burlington, MA, USA |
| GNAT | Mouse | 1:500 |  | Gifted by Jim Hurley, UW, Seattle, USA |
| GFP Polyclonal | Chicken | 1:1000 | # **A10262** | Thermofisher Scientific, Waltham, MA, USA |
| ARCUS-RHO1-2 | Rabbit | 1:1000 |  | Precision BioSciences, Durham, NC, USA |
| Iba-1 | Goat | 1:200 | #**AB5076** | Abcam, Cambridge, MA, USA |
| GFAP | Rabbit | 1:1000 | # **PA316727** | Thermofisher Scientific, Waltham, MA, USA |

| Antibody | Host Species | Working dilution | Cat# | Manufacturer |
| --- | --- | --- | --- | --- |
| **Secondary** |  |  |  |  |
| Alexa Fluor 488 anti-chicken IgG | Donkey | 1:1000 | # **703-545-155** | Jackson Immuno Research Laboratories Inc., Westgroove, PA, USA |
| Alexa Fluor 546 anti-mouse IgG(H+L) | Donkey | 1:1000 | # **A10036** | Thermofisher Scientific, Waltham, MA, USA |
| Alexa Fluor 647 anti-rabbit IgG (H+L) | Donkey | 1:1000 | # **A32795** | Thermofisher Scientific, Waltham, MA, USA |
| Alexa Fluor 568 anti-goat IgG (H+L) | Donkey | 1:1000 | # **A-11057** | Thermofisher Scientific, Waltham, MA, USA |

### Vision testing of pigs

We adapted an obstacle course for pigs(*45*) to define differences in mobility under rod-isolated (0.01 cd·m^-2^) or cone-mediated (20 cd·m^-2^) retinal signaling in age-matched RHO1-2 treated and untreated Tg and in WT pigs. The maze was built from plastic traffic barriers (orange), hay bales (white), and heavy-duty fencing (green). The position of the traffic barriers was randomized (3 configurations), and the distance between barriers varied to prevent memorization of the maze. Pigs were trained to navigate the maze under light-adapted conditions, receiving a food reward at the end. The next day, pigs were fasted overnight, brought into a holding pen, and dark-adapted for 30 mins. Under rod-isolated conditions (experimenters could no longer see color), the pig’s navigation through the maze was recorded by a hand-held camera in IR mode and with IR room illumination. Pigs wore a mask over their nose and face with eye holes that could be covered so that they could be tested either binocularly or monocularly. Pigs were tested several times over a week, with the maze course randomized between runs. Videos were brought into a motion analysis program (Kinovea), and the position of the front feet relative to the obstacles was plotted and calculated using SmartDraw. The variables measured and compared across WT, uninjected and RHO1-2 injected pigs were: collisions, time, path length and steps needed to complete the course.

### Statistical Analyses

Statistical tests were performed for all measures using GraphPad Prism v10.1 software (GraphPad software, Sandiego, CA, USA). All data presented represent Mean ± SEM. Before statistical testing, the normality of the data distribution was determined using the Shapiro-Wilk Test. Statistical significance was determined using the appropriate statistic, e.g., parametric One-way ANOVA test (for normally distributed data) or non-parametric Kruskal-Wallis test (for non-normal data). When the overall ANOVA was significant, multiple comparisons were performed using post-hoc Sidak and Dunn’s multiple comparison tests for parametric and nonparametric tests, respectively. The p-value for significance was set at p<0.05.

## Acknowledgements

We thank Robert V. Brown, Elyse Frye, Maha Jabbar, Olivia Jacobs, Kynan Jarrett, Gary Owens and Sharon Willer for technical support and Karen Powell, DVM, and the UofL Large Animal Veterinary Staff for their help with the animal assessments. We thank the National Swine Resource and Research Center (U42OD011140) for providing swine, NSRRC:0017.

## Funding

This work was funded by grants:

National Institute of Health grant RO1026158 (MAMc, RGG)

Jewish Heritage Foundation (MAMc)

Kentucky Lions Eye Research Endowed Chair (MAMc)

Preston Joye’s Pope Endowed Chair (RGG)

Precision BioSciences (MAMc)

National Institute of Health grant U42OD027090 (KDW, JAG)

## Authors Contributions

AJ helped to conceive the experimental design, performed the ERG, OCT, and IHC experiments, analyzed all the OCT, ERG, and IHC data, wrote, edited and prepared the manuscript for submission. JMY performed all the RHO1-2 meganuclease editing analyses and edited the manuscript. JF developed the software and analyzed the behavioral data and edited the manuscript. WW performed the majority of the subretinal surgeries. FB helped design and perform the behavioral assays. KK helped to collect ERG and OCT data; analyzed the behavioral data and edited the manuscript .CM helped to collect behavior data. JMN helped conceive the experimental design, helped perform ERG, OCT, and IHC experiments, and analyze ERG and IHC data and edited the manuscript. DCA wrote all custom software for data analyses. SN and JCP analyzed the OCT data. GJ analyzed the behavioral data. JWF and BS helped to perform ERG and OCT experiments and edited the manuscript. HJK performed some of the subretinal surgeries and edited the manuscript. JAG helped with research design and edited the manuscript. KDW helped with research design and edited the manuscript. VVB produced RHO1-2A. JL, MD, KSE performed and analyzed rhAmpSeq, oligo capture, and bioinformatics. R van deB performed and analyzed rhAmpSeq, oligo capture, and bioinformatics. JEC helped to conceive the experimental design. WCL – Performed 2G11 cell experiments and dPCR (Fig. 2C). CT Generated 2G11 cells and ran CHO assay (Fig. 2A-B). DJ Identified nuclease target site for strategy and edited the manuscript. KDV helped to conceive the experimental design, helped to collect the mobility data and edited the manuscript. RGG performed all genotyping, oversight of genome editing analyses and wrote and edited the manuscript. JS optimized ARCUS nucleases to inform strategy for RHO1-2A and built RHO1-2B and, wrote and edited the manuscript. MAMc conceived the experimental design, performed ERG, OCT, and IHC analyses, and wrote and edited the manuscript.

## Data and materials availability

All data associated with this study are present in the paper or the Supplementary Materials. Data supporting the findings of this study are provided in the data files. Sequencing data have been deposited and are available on NCBI SRA, bioproject accession IDs PRJNA1332595 and PRJNA1328906.

## List of Supplementary Materials

Table S1

Figs. S1 to S6

---

**Fig. S1.**
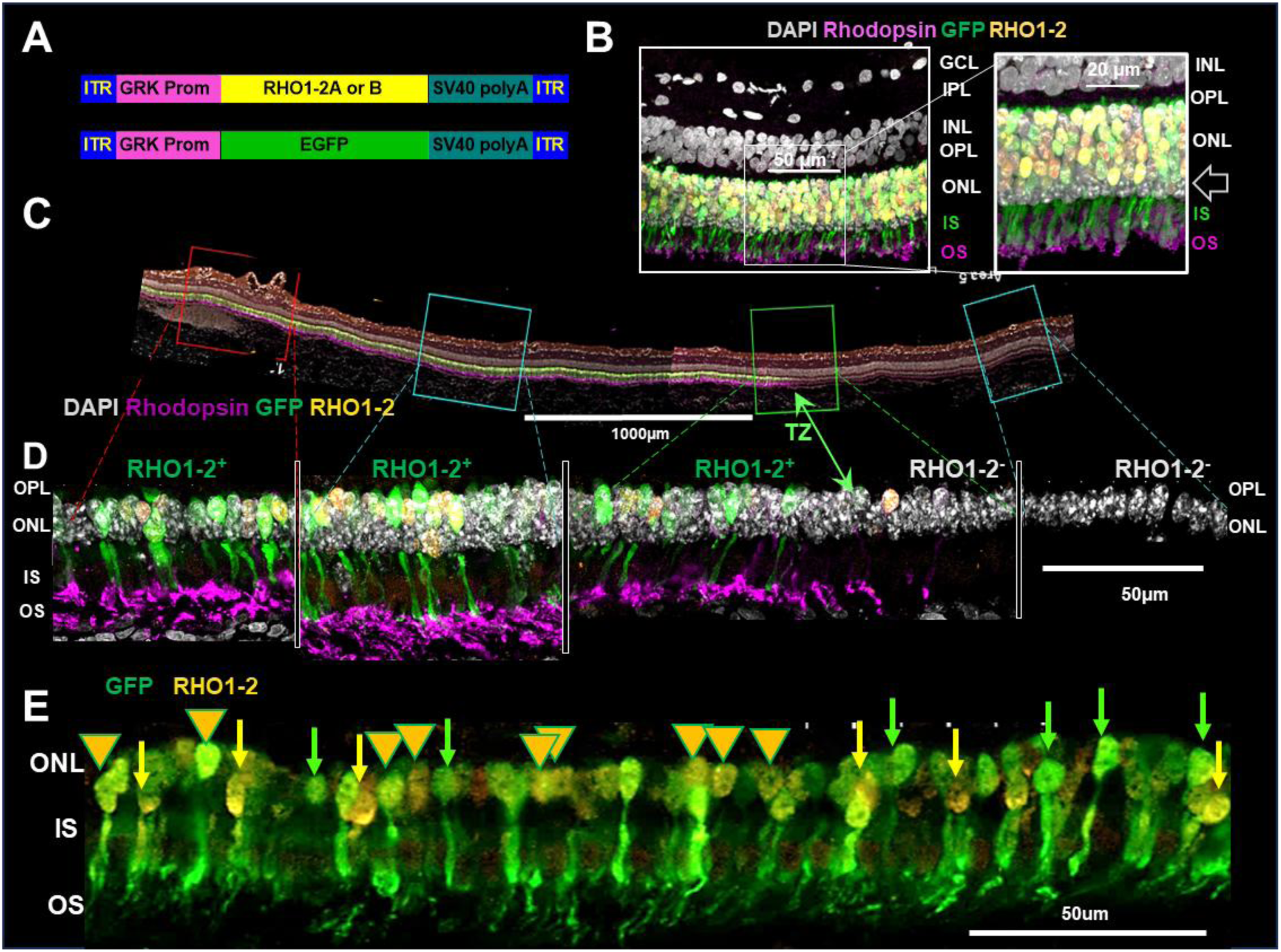
RHO1-2 and GFP expression overlap in rod photoreceptors. (**A**) Schematic diagrams of scAAV:GRK1 vectors: RHO1-A. B and GFP. (**B**) Confocal image of a WT retina subretinally injected with scAAV-GRK1:RHO1-2 + scAAV-GRK1:GFP demonstrating overlapping expression in rods (arrow, uninfected DAPI+ cones). (**C**) Stitched confocal image of a representative retinal section (∼5mm) from a RHO1-2 treated TgP23H pig (2x10^10^vg/eye) showing areas with overlapping expression of RHO1-2 and GFP (RHO1-2^+^), or without expression of either (RHO1-2^-^) and the transition zone (TZ, green arrow). (**D**) Higher power confocal images of individual areas across A (boxes). GFP and RHO1-2 expression ends at the TZ. (**E**) A 250µm long section within a RHO1-2^+^ area in different treated Tg retina illustrating rods expressing both GFP and RHO1-2 (arrowheads), only GFP or only RHO1-2 (green or yellow arrows, respectively). Similar results were found in all retinas (2x10^10^vg/eye; n=11).

**Fig. S2.**
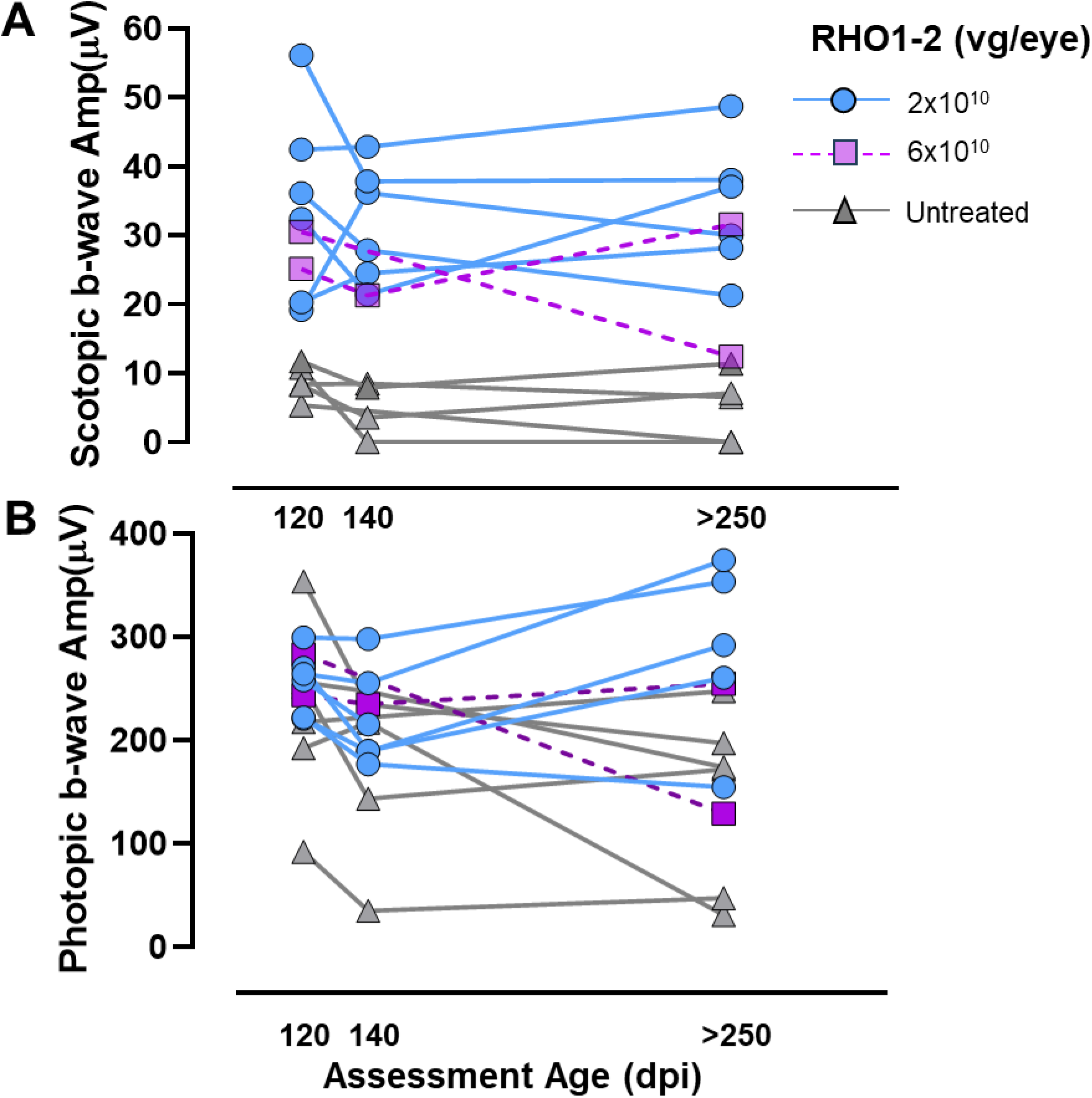
RHO1-2 treatment shows equal efficacy in each TgP23H pig. (**A**) RHO1-2 at 2 and 6x10^10^vg/eye produce similar resurrection of the rod isolated b-wave in TgP23H pigs through ≥250dpi. (**B**) RHO1-2 at 2 and 6x10^10^vg/eye produce consistently higher photopic b-waves in TgP23H pigs beginning at 140dpi.

**Fig. S3.**
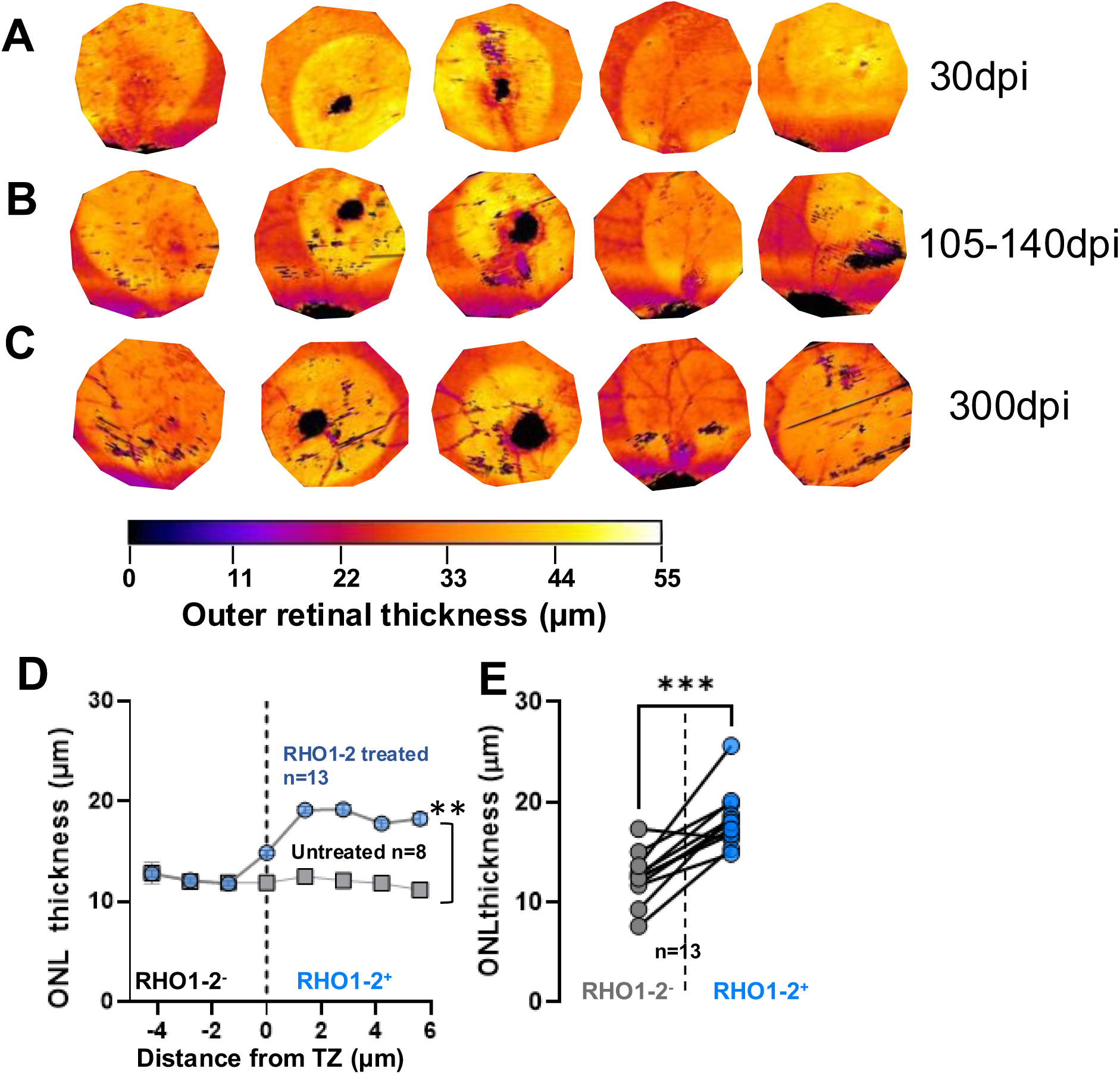
Outer retina and ONL thickness within the RHO1-2+ areas is preserved across time post injection in TgP23H pigs. Heatmaps of the five Tg eyes (columns) at 30 (**A**), 105-140 (**B**) and 300 (**C**) dpi show the stability of the outer retinal thickness (OS/IS + ONL) in RHO1-2+ areas (yellow) across time. In 2/5 eyes there is a small area of damage surrounding the retinotomy (black). (**D**) Quantification of SD-OCT b-scans (Fig. 4) shows the ONL is significantly thicker within RHO1-2+ areas compared to similar locations in untreated eyes (D: p=0.006; unpaired t-test) and to RHO1-2-areas in the same eye (E, p=0.0001; paired t-test).

**Fig. S4.**
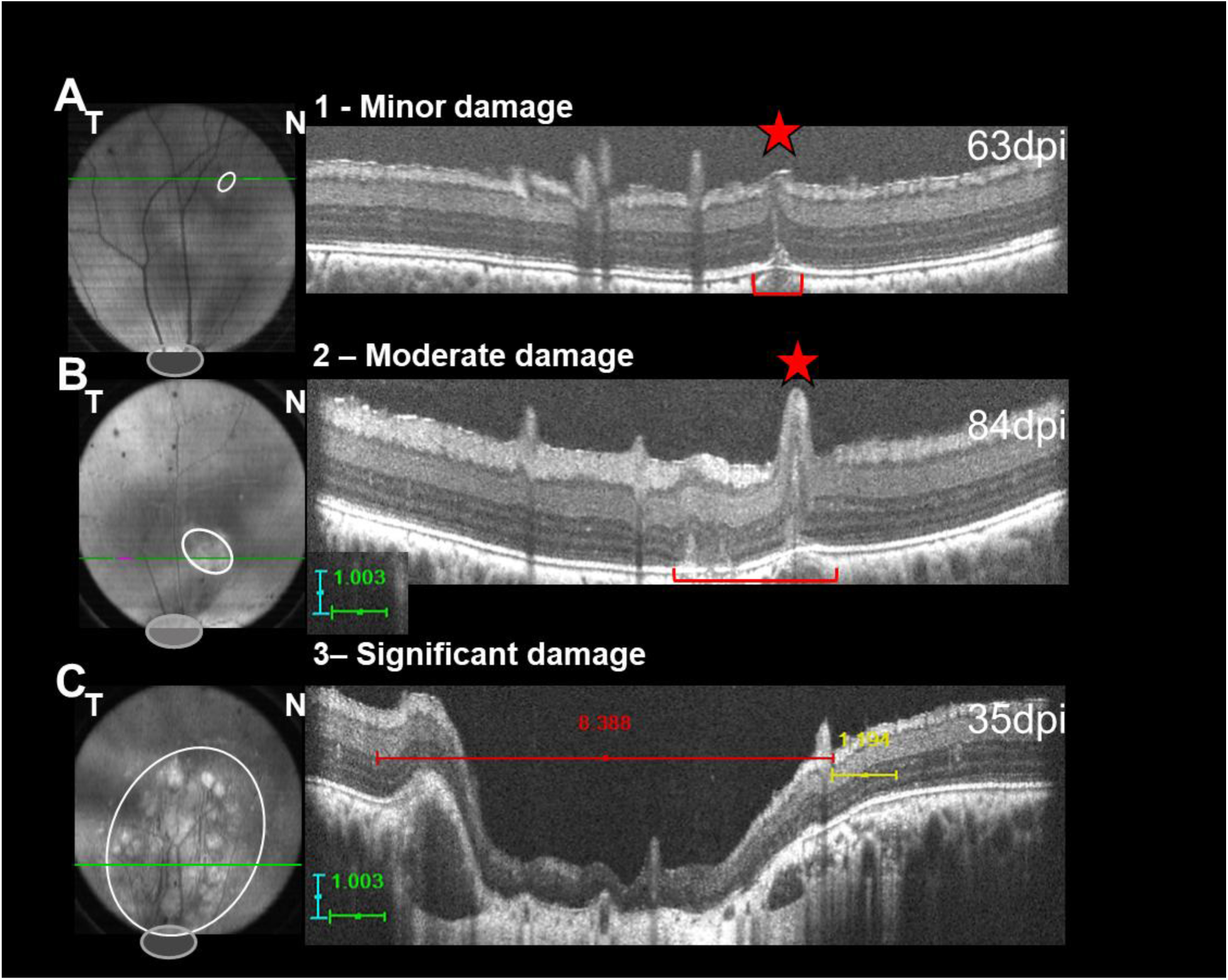
RHO1-2 subretinal injection produce little or no damage to retinal lamination. Representative sdOCT fundus images (right) and b-scans (at the level of the horizontal green line) indicate the site and size of damage (white ellipse, red bracket) caused by subretinal injection (2 x10^10^vg/eye). We qualitatively graded the damage (**A**) 1 – only damage at the site of the retinotomy (star); (**B**) 2 – damage that surrounded the retinotomy that was smaller than the ONH (indicated with the grey ellipse) and only disrupted the outer retina lamination; or (**C**) 3 – damage that disrupted the entire retinal lamination that was larger than the ONH. Category 3 damage occurred only in our earliest injections (25%; 2/8 eyes) or at the same frequency with RHO1-2A (6 x10^10^vg/eye; 2/8 eyes). For all other eyes (n=18; ≤2x10^10^) 89% had damage only at the site of the retinotomy.

**Fig. S5.**
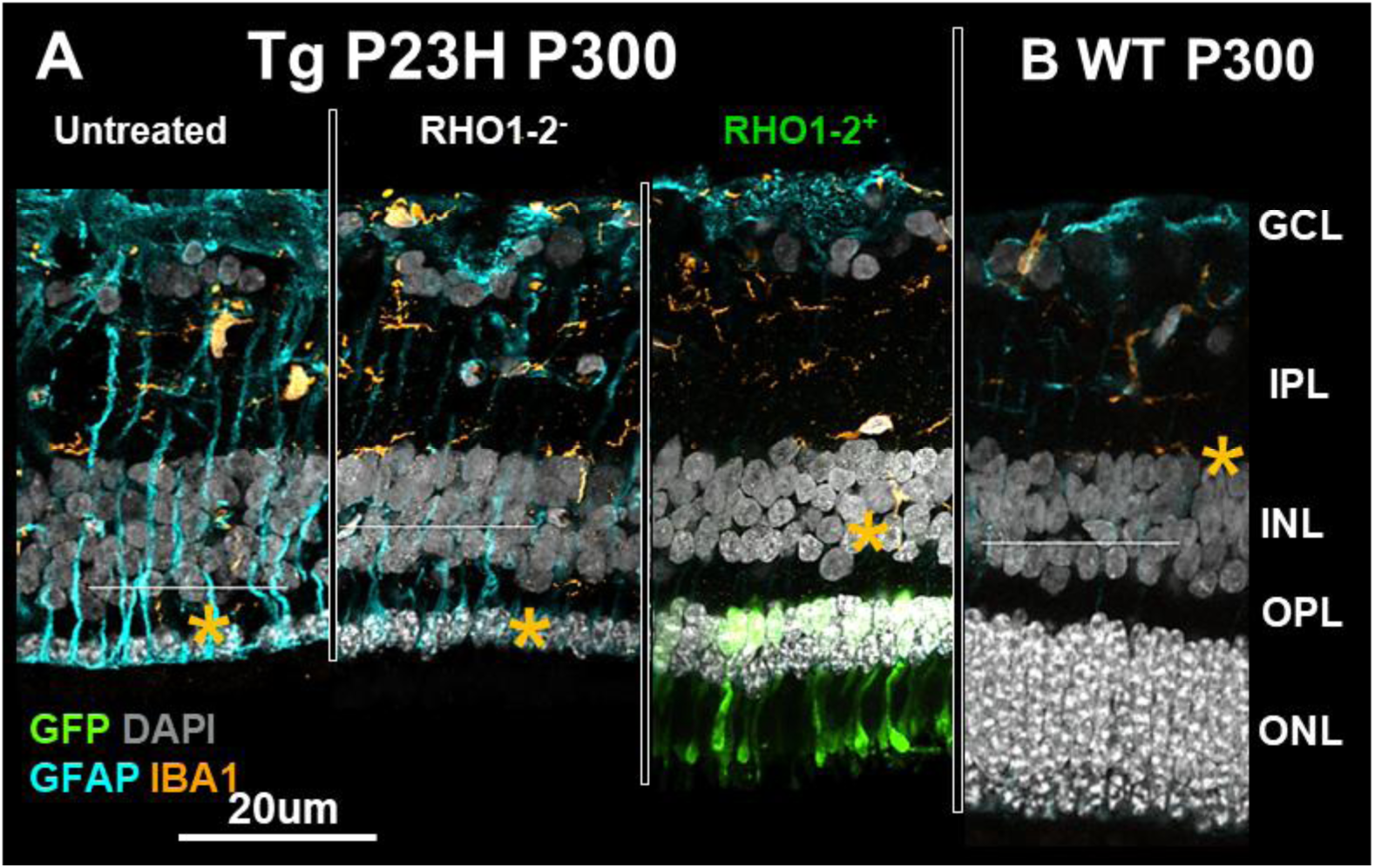
RHO1-2 treatment reduces reactive gliosis in TgP23H RHO1-2+ areas at 140 and 300dpi. (**A**) Confocal images of GFAP+ Mueller Glia and IBA1+ microglia expression in untreated TgP23H pig retina (n=4), in RHO1-2+ and RHO1-2-areas (n=11) and in an age-matched WT pig (**B**). Asterisks (*) show the limits of IBA1+ profiles, which extend to the ONL in untreated (n=4) and RHO1-2-areas (n=11). IBA1+ profiles do not cross the OPL in RHO1-2+ and WT (n=5) retinas. Mueller glia hypertrophy is high in untreated and RHO1-2-areas.

**Fig. S6.**
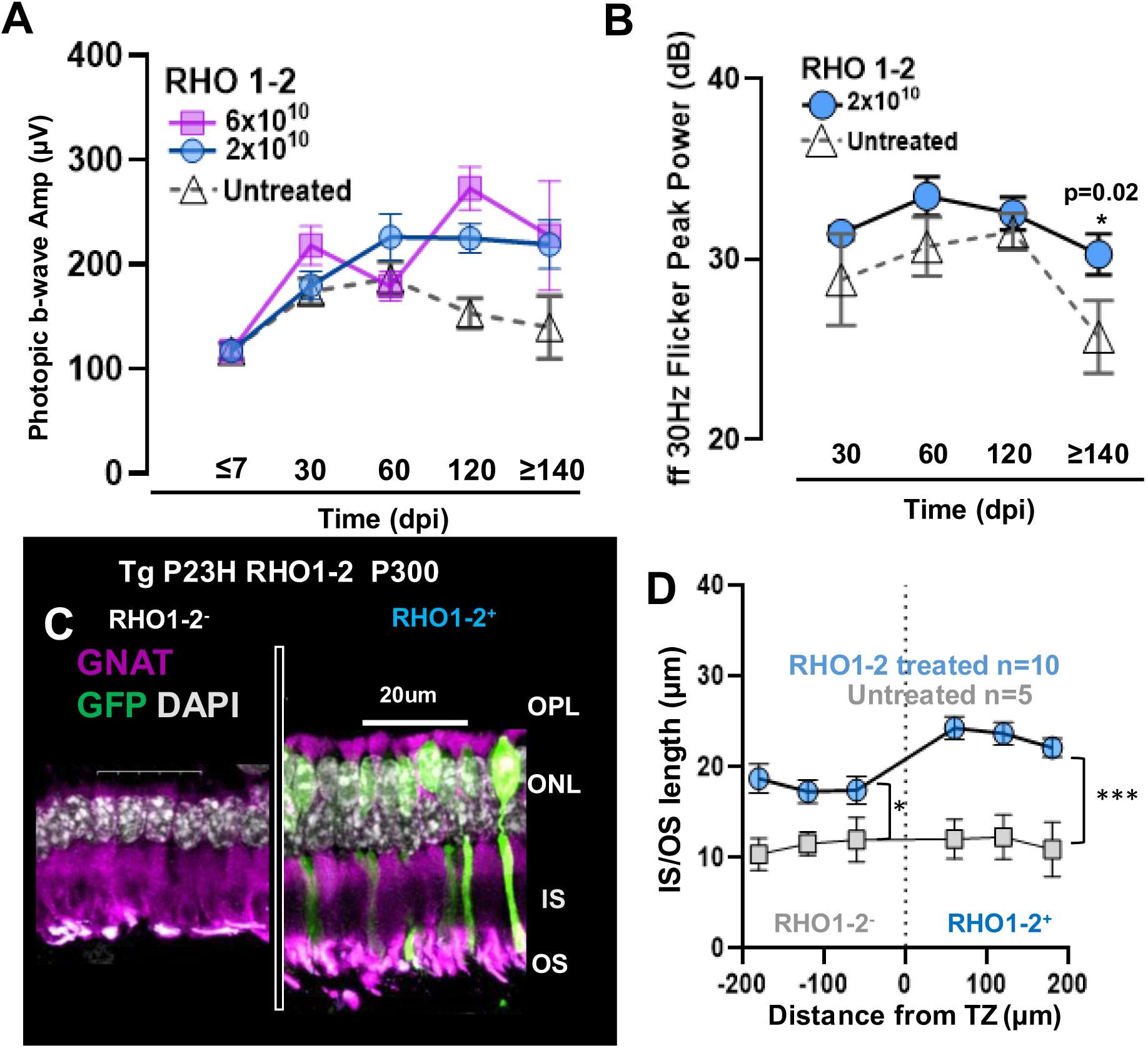
Cone function and structure are improved in RHO1-2 Treated TgP23H retinas. (**A**) Summary of the ffERG photopic b-wave amplitudes as a function of RHO1-2 dose. (**B**) Power of the Flicker ERG (30Hz) in Tg pigs treated with 2x1010vg/eye. Photopic responses indicate that RHO1-2 treatment (P3-7; 2×1010vg/eye) produces significantly larger cone-driven responses compared to untreated at >120dpi (A, B; 2-way ANOVA; Tukey’s multiple comparison test; p<0.04; 60 to >140dpi). The photopic b-wave amplitudes do not reach WT, similar to rod-isolated results. (**C**) Cone morphology is significantly improved in RHO1-2+ areas compared to untreated retina in the same locations. (**D**) Cone IS/OS length in RHO1-2+ is significantly longer than cones in untreated Tg retinas (p=0.0007; Welch’s t-test) and cone IS/OS in RHO1-2-areas are significantly longer than in untreated retinas (p=0.02).

**Table S1.**
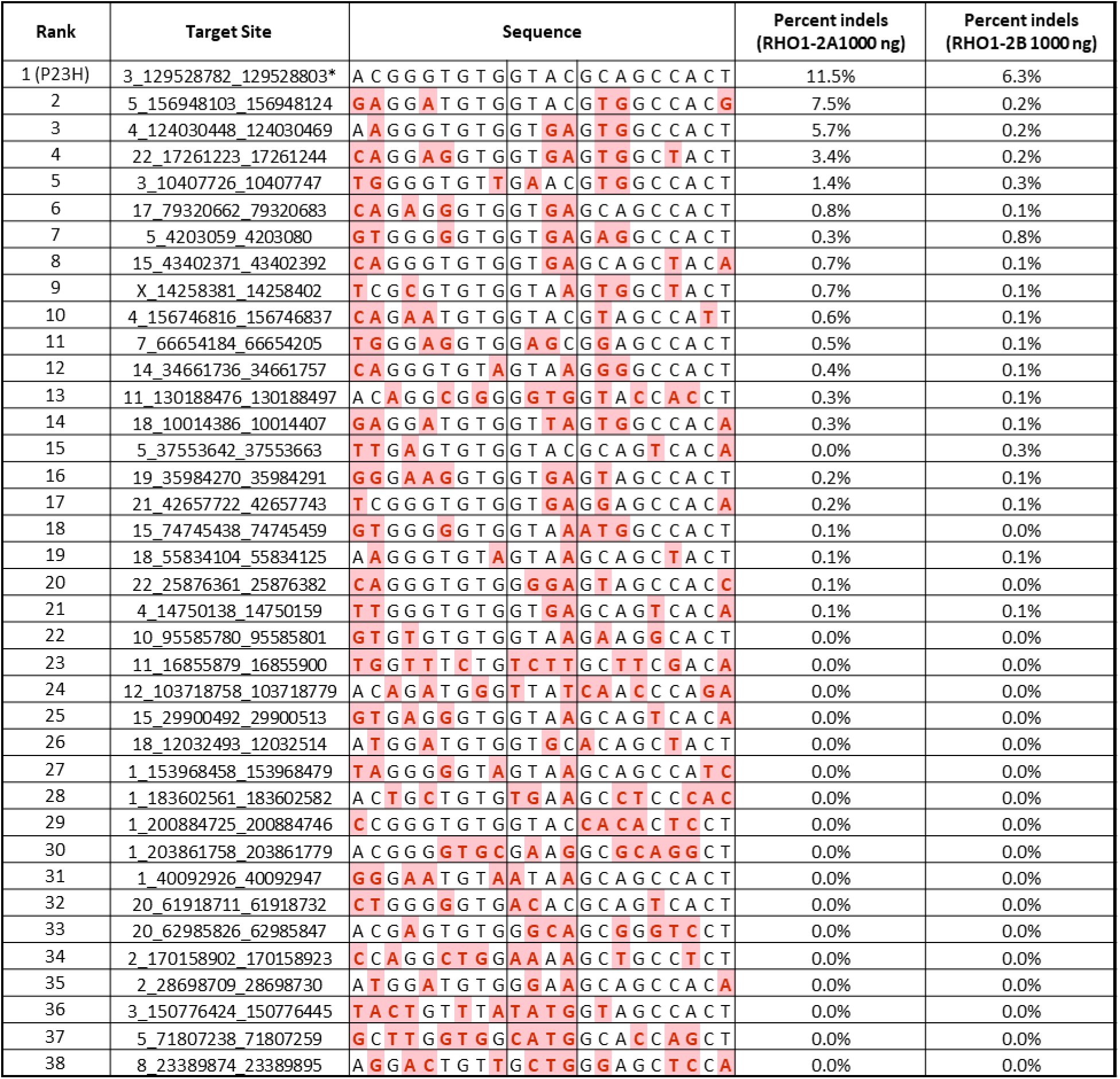
Potential Target sites for RHO 1-2A and RHO 1-2B from nominated sites using NGS.

